# Early Neolithic forest farming at Seven Springs, Martlesham, Suffolk inferred from sedaDNA and pollen

**DOI:** 10.1101/2023.10.11.561859

**Authors:** Samuel M. Hudson, Steven J. Allen, Inger Greve Alsos, Julie Curl, Peter D. Heintzman, Paul Hughes, Lynne Gardiner, Lindsay Lloyd-Smith, Ben Pears, Rob Scaife, Kathleen R. Stoof-Leichsenring, Antony Brown

**Author notes:** Author for correspondence ✉.

## Abstract

The importance of small wetlands and springs to Mesolithic cultures is well established. However, few studies have focused on their significance and use by Early Neolithic agro-pastoralists. Here we present a multiproxy palaeoenvironmental analysis, including ‘authenticated’ sedimentary ancient DNA (sedaDNA), at Seven Springs, Martlesham, UK, indicating that the springs provided an attractive location for pastoral and ritual activity around a palaeochannel surrounded by dense woodland. We demonstrate that sedaDNA can be preserved well within stratigraphically complex wetland sediment sequences, allowing archaeologically valuable insights into crucial periods of change in the prehistoric, where other forms of environmental evidence can be scarce.

## Introduction

Early Neolithic farming in the British Isles has been reassessed in recent years and the arable component shown to be both spatially and temporally variable (Stevens and Fuller 2012; Rowley-Conwy 2011; Rowley-Conwy *et al*. 2020). The extent and nature of forest clearance is critical as is the validity of any forest-farming evidence (Clark 1947; Brown 1997; Innes *et al*. 2013; Fyfe *et al*. 2013; Treasure *et al*. 2019). Were early farmers using small, possibly even pre-existing, clearings and utilizing the surrounding woodlands, rivers, and streams for grazing and watering? Excavations in the parish of Great Bealings, Suffolk, England, in an area near Martlesham known locally as Seven Springs (Figure 1) have revealed an extensive archaeological record which has bearing on this question. Fieldwork was carried out as part of archaeological mitigation for the onshore cable corridor of the East Anglia One Offshore Windfarm. The site, which has been designated East Anglia One Site 23 (parish code BEG 059, Suffolk Historic Environment Record), includes a number of peat-infilled paleochannels, one of which contained large amounts of preserved wood, including some worked timber, along with pottery, struck flint and animal bone dated to the Neolithic and Early Bronze Age. The excellent preservation within this shallow palaeochannel (Palaeochannel 1), is due to the presence of at least two ground-water springs, which maintained saturated conditions.

**Figure 1:**
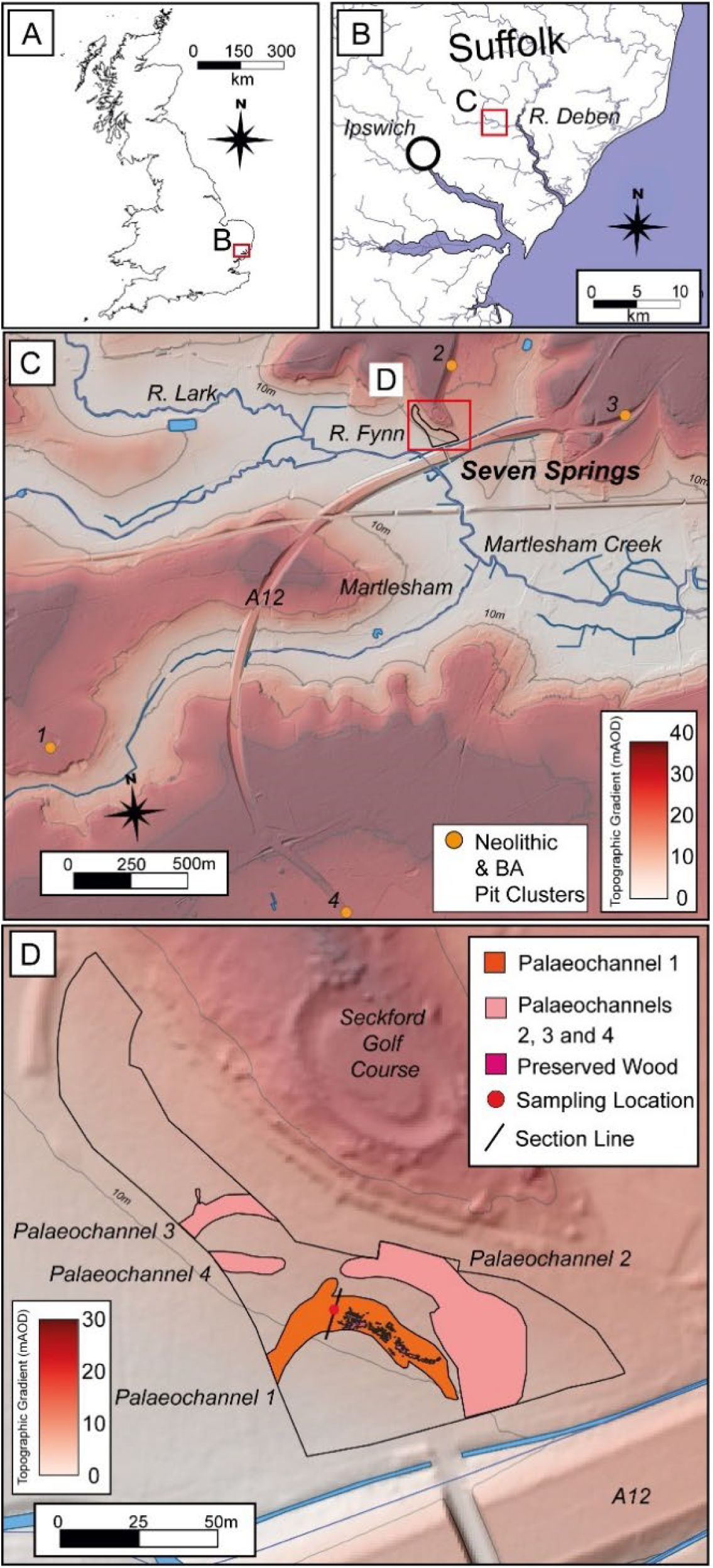
(A) Location of Deben River Catchment, Suffolk, UK. (B) Position of excavation site at the head of the River Deben intertidal zone and confluence with the River Fynn. (C) Position of excavation area on northern edge of the River Fynn. Neolithic and Bronze Age Pit Cluster Sites (Garrow 2006) are as follows: 1- Little Bealings, 2-Great Bealings, 3-Martlesham Bypass, 4- Martlesham Heath. (D) Palaeochannels identified with extent of wood, trench section line and sample location across Palaeochannel 1.

Springs attract both human and animal activity. A concentration of human activity is seen at Late Mesolithic sites such as Cherhill and Blick Mead in Wiltshire (Evans *et al*. 1983; Jacques *et al*. 2018; Hudson *et al* 2022.), Nine Wells in Cambridgeshire (Boreham *et al*. 2020), and Langley’s Lane, Somerset (Lewis *et al*. 2019). In the early Neolithic, springs are found inside causewayed enclosures such as Freston, Suffolk (Carter *et al*. 2022) and are closely associated with many of the large, monumental mounds seen in the Late Neolithic and Bronze Age such as those at Silbury Hill, Marlborough Hill and Hatfield Barrow (Leary *et al*. 2010, 2012), and stone monuments (e.g. Darvill and Wainright 2014). Moreover, it has been argued that the manipulation and incorporation of groundwater, such as the creation of an infilled ditch, was integral to the symbolism of Late Neolithic henge monuments, acting as part of a wider representation of landscape (Richards 1996). Springs can also be the focus of intensive ritualised activity, linked with other freshwater waterbodies and conduits in a wider ritualistic association of water as a bridge between the living and the dead, marked by ritual deposition (Davies and Robb 2004; Bradley *et al*. 2015).

A number of features found at Seven Springs suggest heightened prehistoric significance of the site (Figure 2); including the presence of a carefully deposited or curated aurochs skull (dated 4449-4267 cal. BC) and roe deer antler frontlets with cut marks and areas of worn polish on the interior of the skull, suggesting use as a headdress. The distribution of these artefacts within Palaeochannel 1 near to the ground-water spring suggests that the springhead was likely the main reason for archaeological occupation and use of the site. The obvious comparison for the roe deer headdresses is Star Carr, where six were found although with an earlier date at c. 9000 BC (Elliott *et al*. 2019) and others are known from the Mesolithic site of Bedburg-Königshoven in northern Germany (Wild *et al*. 2020). The deposition of the aurochs skull close to the springhead is similar to the Late Mesolithic site of Blick Mead, Wiltshire (Jacques *et al*. 2018), where numerous aurochs bones are found near springs. Thus, the site has both Mesolithic elements and Early Neolithic elements with Neolithic dates (Figure 3) raising the question as to the nature of the site in both subsistence and cultural terms.

**Figure 2.**
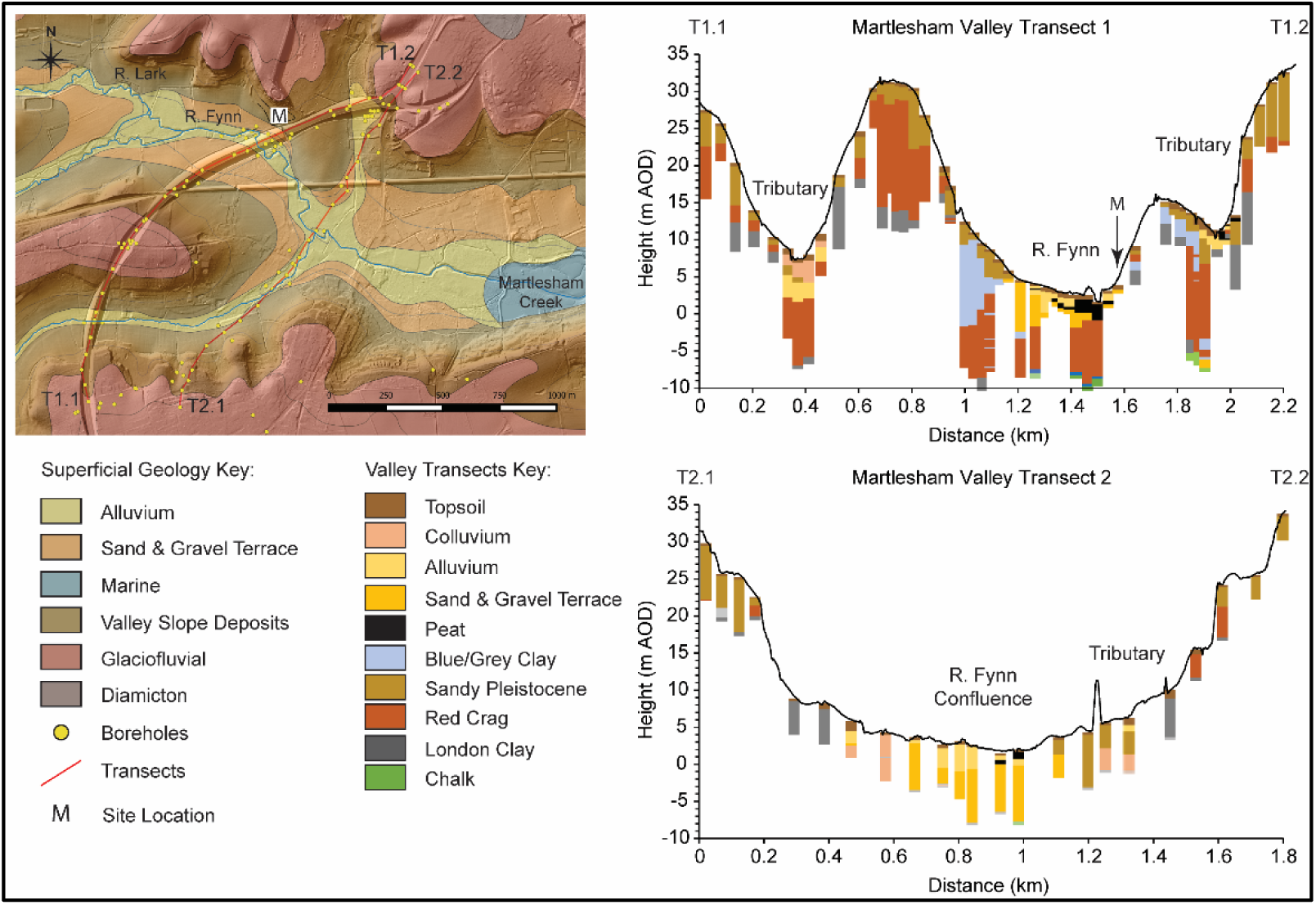
The superficial geology of Martlesham Valley from two transects undertaken by the British Geological Survey (BGS). Constructed in QGIS from BGS open-source data.

**Figure 3:**
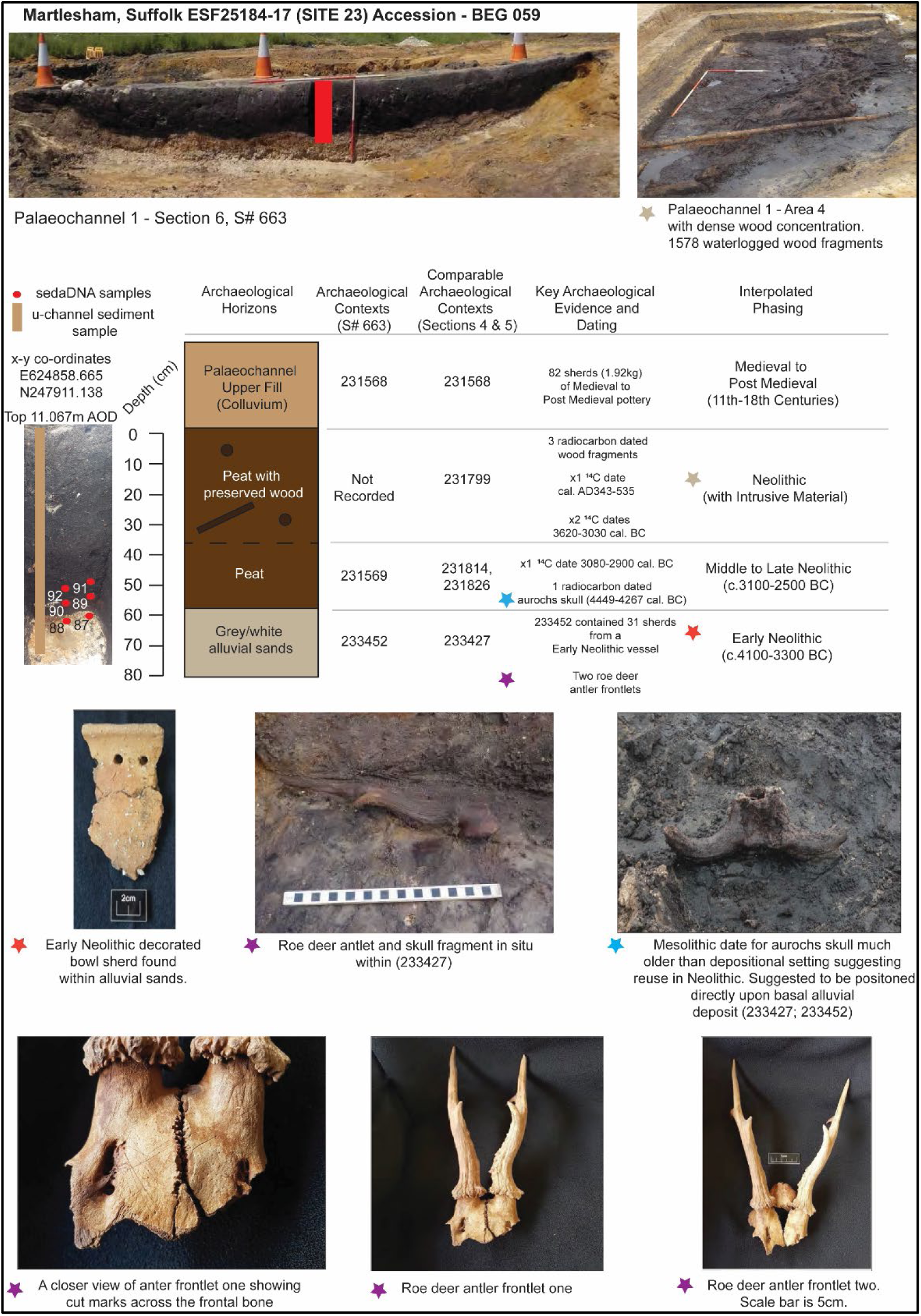
Palaeochannel 1, sampling area 1, section 663, showing the position of the sedaDNA, pollen, and sedimentological samples along with the stratigraphy and key archaeological finds. The dashed line within the stratigraphic diagram illustrates that the boundary between the two peats (231569, 231799) was not clear within the section as no preserved wood was recorded here, Image credits: Archaeological Solutions Ltd.

**Figure 5:**
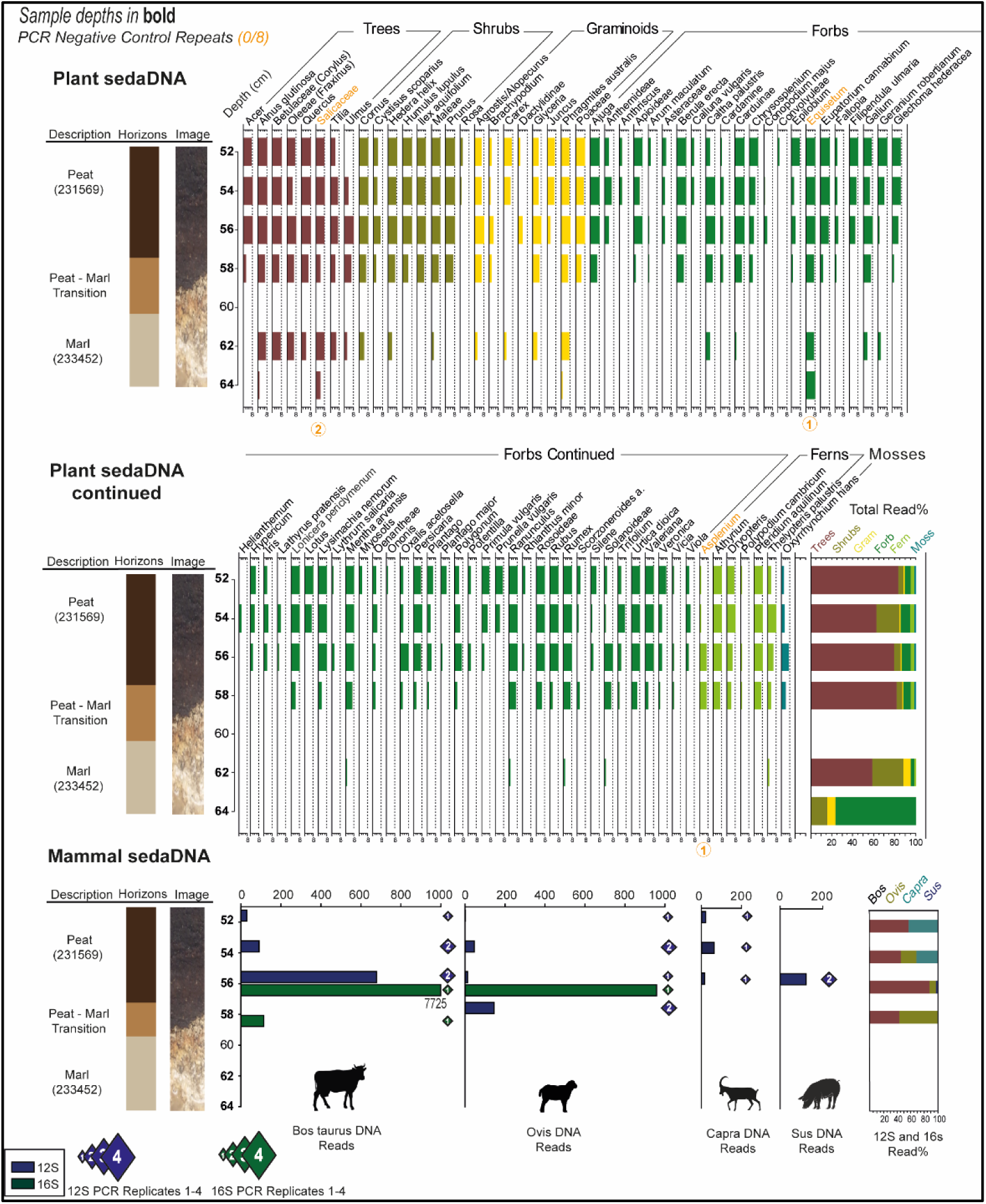
The plant sedaDNA full assemblage from contexts 233452 and 231569 displayed as a number of PCR replicates 1-8 with a composite of total read % on the right. Also shown is the mammal sedaDNA data displayed as combined read and replicate totals from the 12S and 16S markers.

### The Site

The excavated area lies immediately south of Seckford Hall Golf Course, Great Bealings, Martlesham, Suffolk just north of the River Fynn (Long. 1.280463°, Lat. 52.083854°, Figure 1). Initial post-excavation assessment of the site has been completed and a programme of archaeological analysis has begun, commissioned by Scottish Power Renewables as onshore archaeology phase 2 (post-excavation). This paper has been prepared with assessment-level data, as full analysis of the site will not be completed for a number of years, and here only the palaeoenvironmental evidence and Early Prehistoric aspects of the site will be focused on. The site extends along the base and lower edge of the south-facing valley edge slope of the Fynn valley. The site geology (Figure 2) is clays and sands (Thanet Formation), and clays silts and sands (Thames Group) capped to the north by Red Crags Formations sands (Hamblin *et al*. 1997).

Overlying these bedrock strata are three superficial Quaternary deposits; undifferentiated river terrace sands and gravels, Kesgrave Catchment Subgroup fluvial sand and gravels, and, a short distance to the north, an expanse of the glaciogenic Lowestoft Formation sand and gravels. Incised into the superficial geology are a series of palaeochannels infilled with peats and wood and kept saturated by springs issuing from the interface between the Thames Group, Red Crags, and overlying superficial sand and gravel deposits.

### Archaeological Context

The base of the excavated sequence is comprised of alluvial grey sands (233452) which were dated to the Early Neolithic by the ceramic evidence, which included 31 sherds of a decorated bowl (Figure 3) characteristic of the period and of the type (Mildenhall Ware) seen at the nearby sites of Freston, Kilverstone and Mildenhall (Garrow *et al*. 2005; Clark *et al*. 1960:232; Carter *et al*. 2022). These sands also contained 1262g of animal bone, including the deer antler headdresses with their associated skull fragments (Figure 3), which displayed evidence of butchery and polish suggestive of frequent human handling. Further remains from domesticates (cow, sheep/goat) alongside wild species including red deer, roe deer and wild boar were also found on site, though only cattle remains were present in the described section. Above the sands, the lower peat infill (231569) contained 2 other Early Neolithic sherds, along with three intrusive sherds of Iron Age date. Also found at the base of this lower peat infill was an aurochs skull with horn core (4450-4260 cal. BC) that showed cut-marks. The Late Mesolithic radiocarbon date of the skull (Table S10) indicates that the skull may have been reused in the Neolithic and placed deliberately onto the basal alluvial grey sands, but this is uncertain. A radiocarbon dated piece of wood from this layer gave a date of 3080-2900 cal. BC, confirming a Neolithic context.

The overlying peat (231799) contained 1578 pieces of preserved wood, these were not recorded in the analysed section but prevalent across the majority of Palaeochannel 1. 22 pieces were deemed to be worked or possibly worked, most with a metal axe and these pieces are thought to be medieval in date and driven down from higher up. Three pieces of the unworked wood from this layer were radiocarbon dated, with two producing a combined calibrated date range of 3620-3030 cal. BC. and the third a date of cal. AD 340-540, suggesting it is intrusive from the layers above. Overlying this layer, the upper fill of the palaeochannel was a sandy colluvium that contained medieval and post-medieval pottery. In summary, the upper peat with preserved wood horizon (231799) might be disturbed due to root growth within the peat, but the combined radiocarbon and relative archaeological dating suggest that the lower analysed sections of the sequence (64-48cm, (231569), (233452)) are less affected and span much of the Neolithic period (4000-2900 BC). However, there remains a likelihood of disturbance and intrusive material within these contexts, in part due to the complexities of valley floor stratigraphy highlighted by Howard *et al*. (2009). This made further authentication of the DNA evidence through shotgun sequencing and damage pattern analysis necessary (see Results).

## Methods

Environmental analyses were conducted on subsamples from six 50ml falcon tubes (*seda*DNA; Figure 3) and a 70cm u-channel for pollen, loss-on-ignition (LOI), and portable XRF (pXRF) taken from the basal grey sands and overlying peat from excavation area 1, section line 6. *Seda*DNA analysis consisted of both shotgun metagenomics and plant and animal metabarcoding of extracted DNA from the archaeological sediments. Full methodology can be found in Text S1.

## Results

### Sedimentology and pollen analysis

The LOI and pXRF analyses defined the major stratigraphic units (Figure S1). The basal grey sands (233452) described in the field were found to have a high carbonate content (up to 40%) and have been reclassified as a sandy-marl. The pollen assemblage (Figure 6) is dominated by deciduous woodland taxa, with limited herbs present throughout (max. ∼6%). Most notable is lime (*Tilia* cf. *cordata*) with high pollen frequencies in the lower profile and a distinct decline at *c.*56cm. Given the marked under-representation of lime in pollen assemblages, due to a number of factors, the values here suggest it was dominant on drier ground immediately adjacent to the site as was common in the mid-Holocene throughout lowland Britain (Greig 1982; Brown 1988). Oak (*Quercus*), elm (*Ulmus*) and holly (*Ilex*) were also present. Subsequently, there was an increase in hazel (*Corylus avellana*), possibly scrub colonisation after the reduction in elm and lime. Given the geology and topography, this rich mix is due to the close proximity of the palaeochannel to the floodplain edge (*c.* 30m) with pollen coming from both wet and dry woodland communities. The herbs give little indication of any open areas with an unusually low level of grasses (Poaceae), only a trace of bracken (*Pteridium aquilinum*) and few other light-demanding taxa, but high values of the shade-tolerant ferns of *Dryopteris.* Cereal-type pollen of *Avena-Triticum* t. is present (1-2%) suggesting local arable activity. It should be noted that the density of the adjacent woodland will have inhibited pollen input of such possibly localised areas of cultivation to the fen mire. Indications of marsh/wetland are shown by a consistent presence of sedges (Cyperaceae) throughout (1-5%). These herbs will have formed the field layer to the on-site alder-carr floodplain woodland. In summary, the pollen suggests that during the Neolithic the site was surrounded by closed-canopy mixed deciduous woodland.

**Figure 6.**
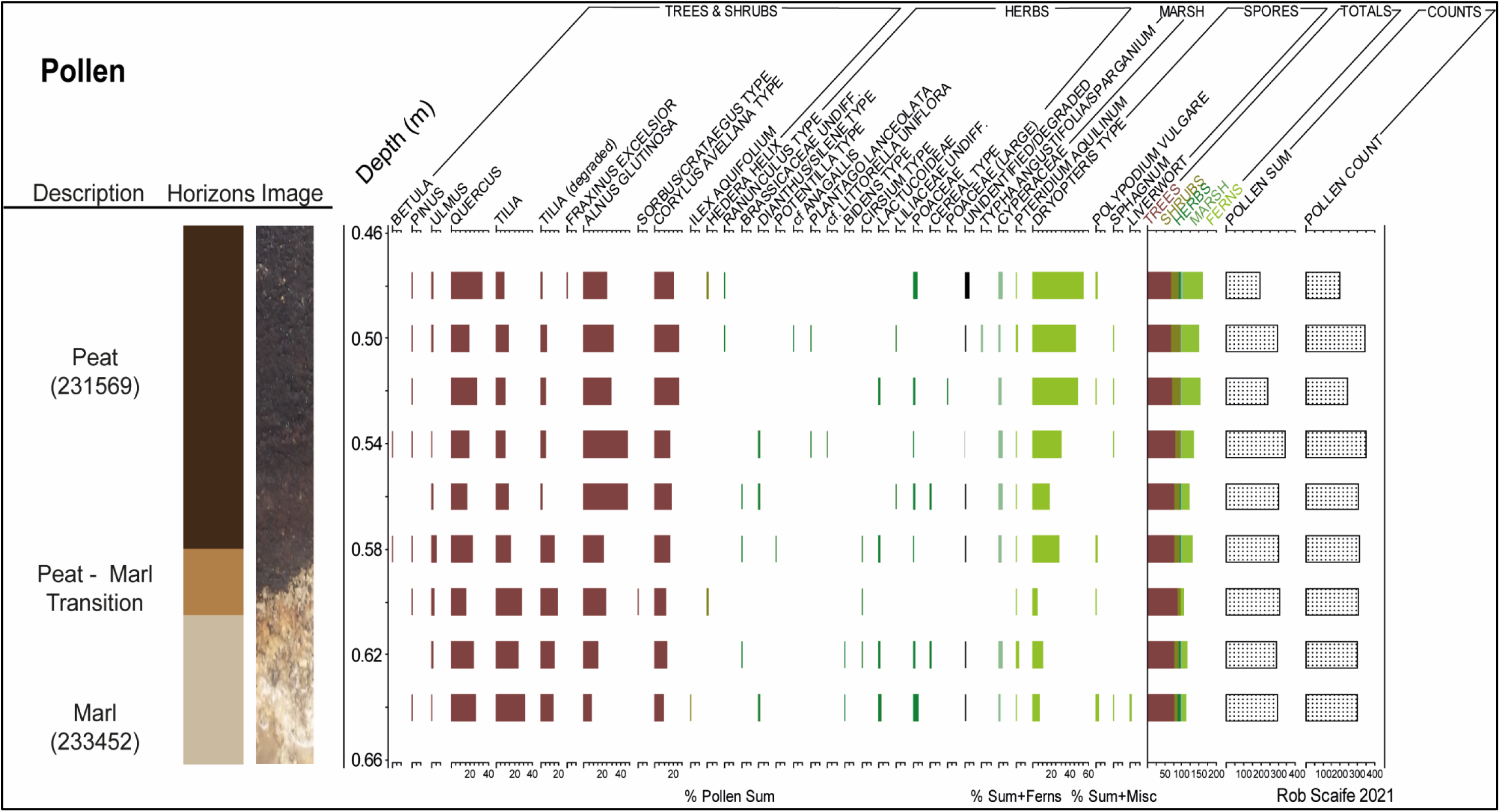
The pollen assemblage, produced from subsamples every 2cm between 48-62cm of the u-channel. The assemblage was dominated by arboreal pollen with a very limited number of herbs. Pollen sum includes total dry land pollen including Alnus and Salix.

**Figure 7:**
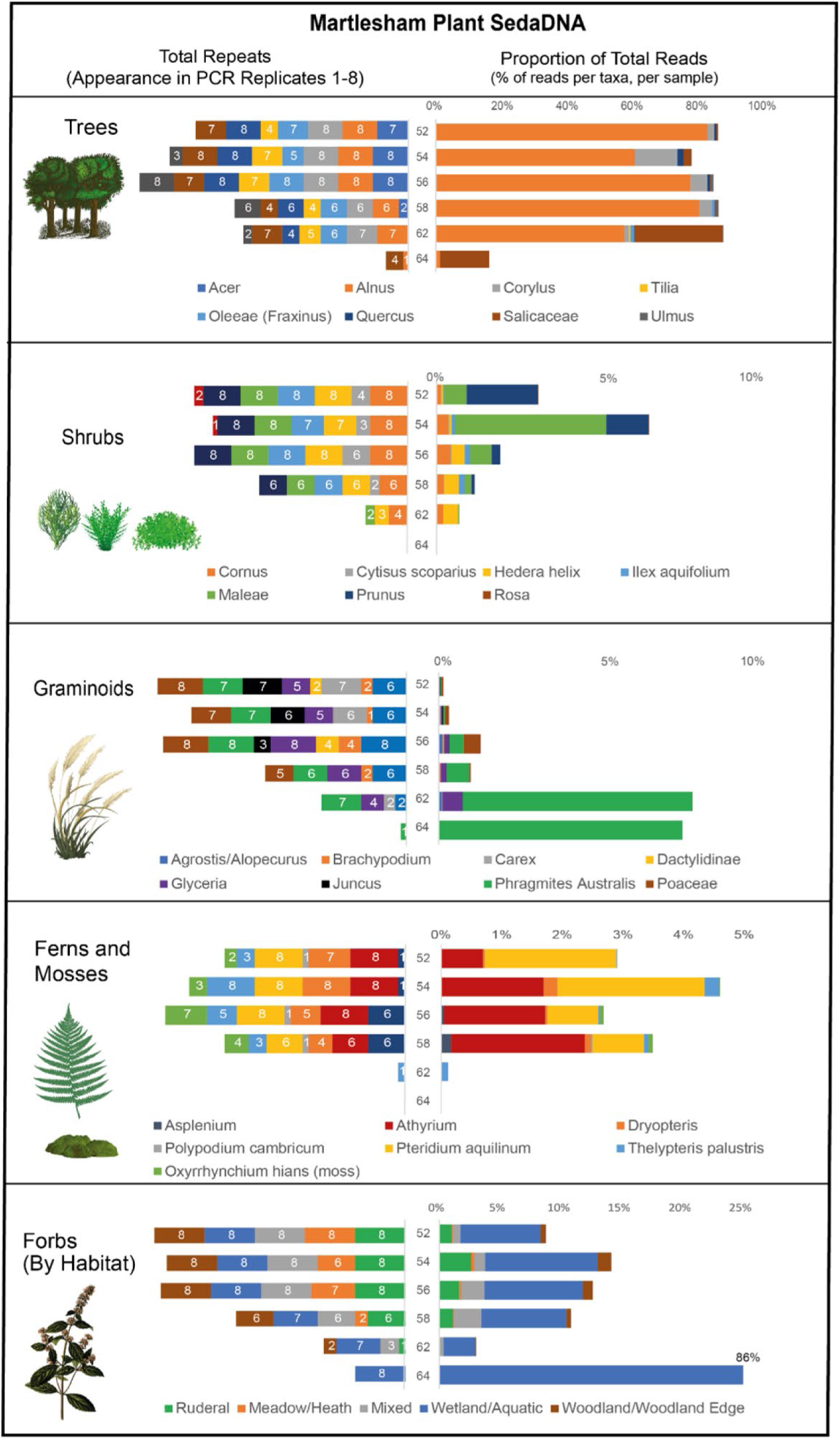
The plant sedaDNA metabarcoding data from Palaeochannel 1 displayed through both number of PCR replicates 1-8 and % of total reads per sample.

### S*eda*DNA Metabarcoding

#### Early Neolithic Marl (233452) (64-62cm)

The two basal *seda*DNA samples taken from within the marl at 64 and 62cm yielded little data, probably due to a low amount of template DNA present. All plant taxa types are seen in largely diminished numbers of PCR replicates and reads, particularly forbs. The sample from 64cm contains only alder, willow (Salicaceae), grasses (Poaceae) and horsetail (*Equisetum*). Within the sample at 62cm the trees consist of high amounts of alder, lime and oak, and some elm. Shrubs and graminoids are seen in slightly lower numbers of PCR replicates and reads, with the exception of the common reed (*Phragmites australis*) which is well represented (8% total reads, 7/8 PCR replicates). Forbs are limited to a range of damp meadow herbs such as marsh marigold (*Caltha palustris*), horsetail (*Equisetum*), herb robert (*Geranium robertianum*) and bedstraw (*Galium*). The marsh-fern (*Thelypteris palustris*) is also seen in a small number of PCR replicates and reads.

#### Neolithic Peat (231569) (58-52cm)

The four upper *seda*DNA samples are largely similar in their composition. Four mammal taxa are seen only in these samples with *Bos taurus* being the most prevalent. The sample at 56cm produced the highest number of reads, mostly of *Bos* and *Ovis*. All four samples contained a diverse community of all types of identified plant taxa, with the exception of mosses, which are represented solely by Swartz’s Feather-moss (*Oxyrrhynchium hians*) that is characteristic of calcicole woodland. Tree and shrub taxa expand to include maple (*Acer*), broom (*Cytisus scoparius*), holly (*Ilex aquifolium*), cherry-sloe (*Prunus*), with the addition of aspen (*Populus*) and rose (*Rosa*) in the samples at 52cm and 54cm. Alder overwhelmingly dominates overall read percentages for these top four samples, probably due to alder wood surrounding the site. Graminoids are augmented by false brome (*Brachypodium*), and common rush *Juncus.* However, marsh taxa such as *Juncus* and *Phragmites* gradually decrease in reads in these samples, suggesting a transition away from river channel vegetation. The forb assemblage is extensive and contains many taxa typical of damp/wet, herb-rich deciduous woodland such as bugle *(Ajuga*), chervils/cow parsley (*Anthriscus*), cuckoo-pint (*Arum maculatum*), meadowsweet (*Filipendula ulmaria*), herb robert (*Geranium robertianum*), ivy (*Hedera helix*), honeysuckle (*Lonicera periclymenum*), yellow pimpernel (*Lysimachia nemorum*) and sorrel (*Oxalis*). Additionally, many taxa found in wetland/marsh habitats such as water-parsnip (*Berula erecta*), marsh marigold, saxifrage (*Chrysosplenium*), willowherb (*Epilobium*), horsetail, hemp agrimony (*Eupatorium cannabinum*), Iris, purple loosestrife (*Lythrum salicaria*) and pale smartweed (*Persicaria lapathifolia*) are present and seen in large proportions of reads (5-15% total reads). Disturbance is also indicated by taxa such as thistles (*Carduinae*), common stinging nettle (*Urtica dioica*), plantain and greater plantain (*Plantago* and *Plantago major*), willowherb, and docks/sorrel (*Rumex*). In the upper two samples, forbs more often found in open environments, such as rock-rose (*Helianthemum*), yellow rattle (*Rhinanthus minor*), trefoil (*Lotus*), meadow vetchling (*Lathyrus pratensis*) and heather (*Calluna vulgaris*), are seen in small read proportions (1-2%). Ferns are well represented, and taxa such as wood-fern (*Dryopteris*) and marsh-fern indicate wet, deciduous woodland. Interestingly, bracken (*Pteridium aquilinum*), which is well represented in PCR replicates in all of the upper four depths, also increases markedly in read proportions in the upper two samples (to 3-4% of total reads).

#### Ancient DNA Damage Pattern Analysis

Aliquots of *seda*DNA extracts from two samples, 52 and 54cm, were used for shotgun metagenomics analysis to (1) test the metabarcoding detections of *Bos* and *Ovis* in these two samples, (2) clarify whether *Bos* is best represented by wild aurochs (*Bos primigenius*) or domestic cattle (*Bos taurus*), and (3) determine whether the Betulaceae taxon identified in metabarcoding data was *Corylus* or *Betula*. We used read mapping statistics and ancient DNA damage-pattern analysis to determine the best supported of these hypotheses (Figures S2-S7; for methods and detailed results, see Text S1 and S2). The shotgun metagenomics data and damage-pattern analyses confirmed the pattern of presence of *Bos* and *Ovis* found within the 12S and 16S datasets and confirmed the presence of ancient DNA derived damage patterns for both the plant (*Betula*, *Corylus*) and mammal DNA (*Bos*, *Ovis*) analysed. Whilst damage patterns cannot ‘date’ the DNA, the damage pattern frequencies of the *Corylus* and *Betula* (0.12-0.19) are high enough to likely be prehistoric as gauged by lake sequences (Brown subm.). Therefore, contamination from modern intrusive material is unlikely to be a significant component of the DNA. Comparison of the *Bos seda*DNA to selected *Bos* reference mitochondrial genomes (Table S9), indicate that the *Bos seda*DNA originated from domestic cattle (*Bos taurus)* as opposed to aurochs (*Bos primigenius*), ruling out DNA recovery from the aurochs skull as a potential source of mammal DNA. Lastly, we find that the Betulaceae taxon is best represented by *Corylus* (hazel), thereby expanding upon and clarifying the identification based on metabarcoding.

#### Environmental Synthesis

The combined lithostratigraphy, *seda*DNA and pollen assemblages identified at Martlesham indicate a palaeochannel that underwent hydroseral succession beginning in the Neolithic after being cut off from the main river channel(s). A small stream-fed pond was then present during the Early Neolithic, as indicated by the Early Neolithic pottery sherds in the marl, before eventual peat development and terrestrialisation. During peat development, the pollen and *seda*DNA data indicate the presence of species-rich deciduous woodland, which is corroborated by the findings of oak, elm, alder, and ash in the preserved wood (Text S3). This woodland was immediately adjacent to the channel and dominated by lime, with alder within the palaeochannel. After lime decreases, hazel and oak become dominant. The *seda*DNA reveals that the site was used by domesticated stock, mostly cattle and sheep but also possibly goat.

## Discussion

The mid-Holocene environment at Seven Springs is typical of low-energy forested systems in the early-middle Holocene (Brown *et al*. 2021), and particularly the East Anglian region (Smith *et al*. 1989; Hopla *et al*. 2009; Geary *et al*. 2016). Floodplain peat development occurred in a palaeochannel followed by succession to alder dominated fen-carr. Previous sedimentological research at Rendlesham (Figure 8) in the Deben valley has indicated that infilling of a main palaeochannel of the River Deben occurred at 4425-4371 cal. BC, with repeated fluvial activation and organo-mineral accretion up to 1065-1058 cal. BC in the Late Bronze Age (French and Taylor 2016:20).

**Figure 8.**
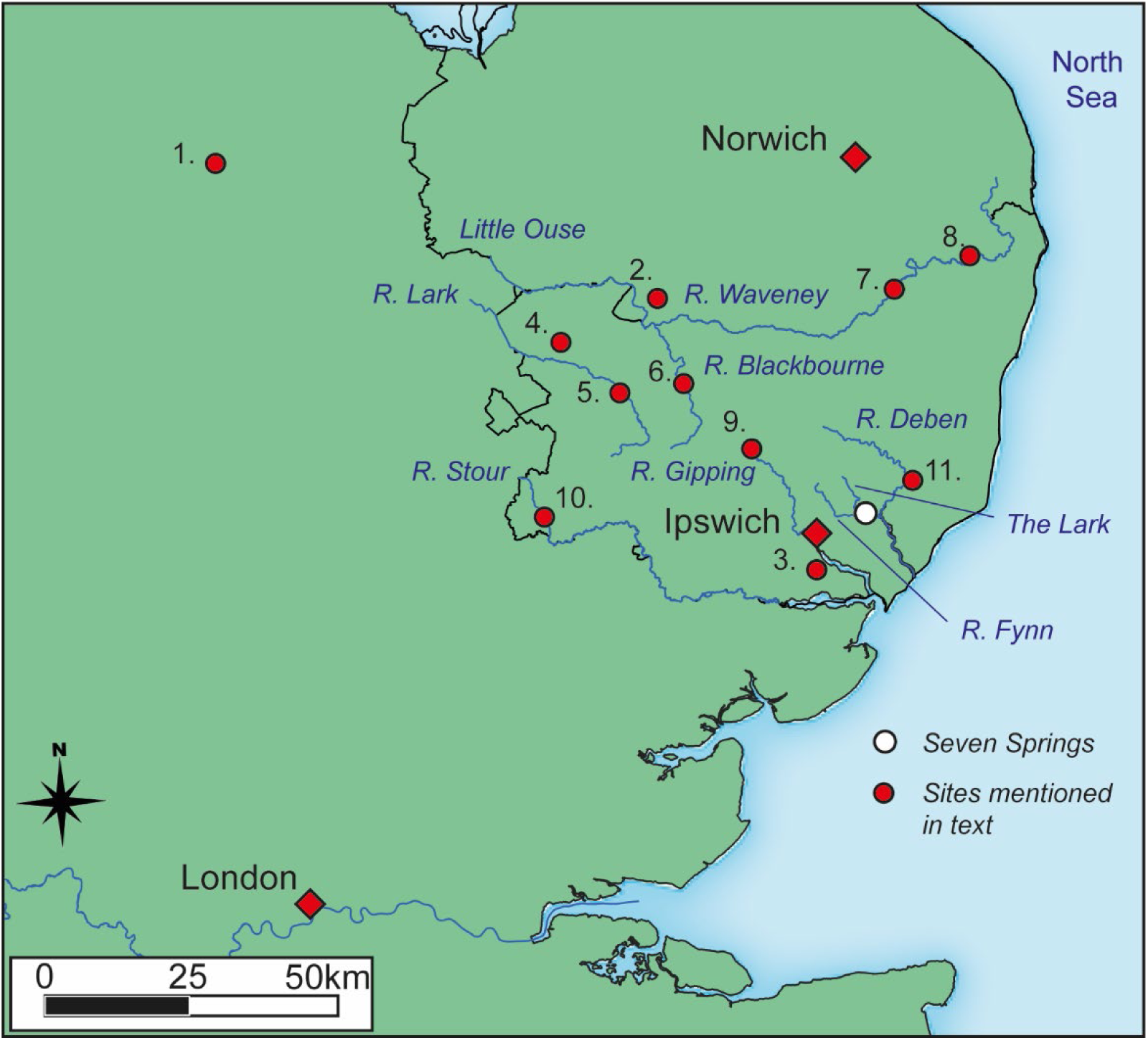
Sites mentioned in text. 1. - Etton, 2. - Kilverstone, 3. - Freston, 4. - Mildenhall, 5. - Fornham All Saints, 6. - Ixworth, 7. -Flixton, 8. - Beccles, 9. - Stowmarket, 10. - Kedington, 11. - Rendlesham.

Comparable environmental sequences can be found in other Suffolk River Valleys, namely those of the Waveney, Lark, Little Ouse, Blackbourne, Gipping and Stour (Geary *et al*. 2016). The *seda*DNA and pollen data from Seven Springs is consistent with pollen sequences from Beccles and Ixworth, in that all suggest that lime was a dominant component of Mid-Holocene dry woodland in the region, with lesser components of oak, hazel and elm (Geary *et al*. 2016). The mid-Holocene carr-woodland dominance of alder in the region is also seen on the River Gipping floodplain at Stowmarket (Hopla *et al*. 2009).

There is evidence for anthropogenic impact on woodland at Seven Springs, a decline in elm was seen in *seda*DNA reads and PCR replicates as well as in the pollen. The onset of the elm decline varies in East Anglia between 4000 – 3000BC (Waller 1994a). More noticeable is a decline in lime pollen above the basal marl, possibly due to the on-site paludification which has been demonstrated to reduce *Tilia* pollen values (Waller 1994b) and aligns with the corresponding increases in alder and hazel pollen in the sequence. Within the surrounding wetland there may also have been slight human impact on lime and elm woodland due to coppicing to create fodder and browse for grazing of the floodplain vegetation. Evidence of coppiced stools are known in the Neolithic for the region, being found *in situ* at Etton (Pryor 2014) and one elm post was found within context 231799 that showed possible working marks with a stone axe (Text S3). There are also some indications of grazing from the variety of woodland edge and ruderal plant species seen in the *seda*DNA data: apple, rose, bugle, bindweed (*Convolvulus*), meadow vetchling, brambles, clover (*Trifolium*), nettle, plantain and bracken (Behre 1986: 233). An alternative explanation for the decreases in *Tlia* and *Ulmus* seen could be a hiatus between the basal sands and peats above.

Woodland exploitation (coppicing, pollarding) and its effects on pollen diagrams has been explored in previous research and remains imperfectly understood (Edwards 1993), but is thought to generally reduce pollen counts of alder and lime over time (Waller *et al*. 2012). The genetic data also suggests that at least three domesticated species were present on site (possibly four if *Sus* represents pig), with cattle dominating the *seda*DNA signature. This is only partially supported by the faunal evidence from the lower peats (Text S4), which shows bones from cattle in Neolithic contexts, but no sheep/goat or pig/wild boar. Together with the cereal grains (1-2%) in the pollen assemblage, this raises the possibility of a nearby clearing for the pasturing of animals seen in the *seda*DNA and faunal assemblages, and some cereal growth. Therefore, one interpretation of Seven Springs is that it represents a site where domestic animals were repeatedly brought to drink at the active springs and graze in the surrounding forest. Nearby, there may have been a settlement or clearing with additional pasture and agriculture. There is some support for Neolithic pastoralism on the floodplain of the adjacent River Gipping, where Geary *et al*. (2016) found dung and dor beetles (earth borers) in Neolithic contexts, suggesting the presence of grazing animals.

In Suffolk, evidence for Neolithic activity is seen in three causewayed enclosures at Fornham All Saints, Kedington and Freston, only the latter having been excavated (Carter *et al*. 2022), along with a recently discovered Early Neolithic long barrow at Flixton (Boulter 2022). More commonly found are pit cluster sites, such as those seen at Hurst Fen and Reydon Farm (Clark *et al*. 1960; Harding 2017), which are common across the whole of East Anglia (Garrow *et al*. 2005; Garrow 2006) and particularly in the Fynn/Deben Valley. Here there are four pit sites located less than a 2km away Seven Springs (Little Bealings, Great Bealings, Martlesham bypass and Martlesham Heath, (Martin and Balkwill 1993), which contained Peterborough ware, Grooved Ware and Beaker type pottery. It is possible that these are near the settlement sites of the people who used and interacted with the springs and palaeochannel complex, establishing it as an area of both functional and ritual use as indicated by the frontlets and aurochs horn-core.

### The Significance of the Springs

The significance of spring sites to Neolithic culture is not often examined in comparison to the much more frequent associations with the Mesolithic. However, the nearby recently excavated causewayed enclosure of Freston, south of Ipswich, provides compelling evidence for the importance of springs to Early Neolithic people (Carter *et al*. 2022). It is hypothesised by Carter *et al*. (2022:97) that the two springs located within the enclosure would have overflowed seasonally, which beside any functional use, created a north-south division that led to spatially structured social actions in the monument, similar to that proposed at Etton (Pryor 2014). In practical terms, the most compelling association with farmers is the easy access to fresh and mineral-rich water for cattle which require around 50 litres per day, and substantially more when used for dairying (Krauß *et al*. 2016).

At Seven Springs, the aurochs skull dated to the late Mesolithic (4449-4267 cal. BC) and was probably deliberately placed into the spring, hinting at the re-use of faunal remains in ritualistic associations with previous Mesolithic cultures, who may have been the first to establish the spring/palaeochannel as an area of importance. This is similar to other spring sites such as Blick Mead, Wiltshire (Jacques *et al*. 2018) and Langley’s Lane, Somerset (Lewis *et al*. 2019). The roe deer headdresses seen in the same context, also have their most obvious counterparts in the Mesolithic, and further remains from both roe deer and red deer were found in the Neolithic contexts. Together the objects suggest a continuity between the two periods, with Neolithic populations using an area associated with the hunting of wild species, for the watering and pasturing of domestic animals seen in the faunal remains and *seda*DNA data. Any significance that the spring held to its previous Mesolithic inhabitants may have been ‘remembered’ through the re-use of the aurochs skull and the creation of the headdresses. In East Anglia as a whole and in particular the fens, settlement patterns inferred from lithic scatters do indicate continuity between the two periods, with Sturt (2006) suggesting that a local understanding of landscape change in the fens may have ‘superseded’ preoccupation over classifications of Neolithic or Mesolithic, hunter-gatherer or farmer. The ritual significance of water and its incorporation into Mesolithic and Neolithic spiritual culture is perhaps unsurprising in East Anglia, an area of the UK where in the mid-Holocene it is likely that sea level change and marine inundation would have been perceptible on a human timescale (Shennan and Horton 2002).

## Conclusions

The sedimentary analysis and dating at Seven Springs revealed a shallow palaeochannel of the River Fynn which formed a spring-fed forest-waterhole in the Neolithic. The *seda*DNA and pollen data showed that the waterhole was within closed canopy mixed deciduous woodland with very little evidence of open land or clearings during the Neolithic. However, mammal *seda*DNA indicated the presence of all four domesticates and particularly cattle which must have been within the small drainage area of the palaeochannel grazing and crossing the floodplain forest. The suggestions of grazing seen in the *seda*DNA and cereals in the pollen pointed to the presence of nearby agriculture/settlement, possibly relating to the surrounding evidence for pit clusters. The data demonstrated how the high taxonomic resolution of authenticated *seda*DNA can provide on-site ecological and resource information above that of pollen as well as the presence of animals and particularly domesticates even from a shallow and stratigraphically complex site. The archaeological evidence of both a curated Mesolithic aurochs skull and roe-deer frontlets suggest cultural continuity with Mesolithic hunting traditions, suggesting awareness of, if not necessarily contact with, hunter-gatherer populations.

## Acknowledgements

Special thanks go to all those involved at Wardell Armstrong with the excavation and post-excavation of the site of Seven Springs, alongside Michael Tetreau, Michael Curnow and Jonathan Tabor at MOLA for assisting with post-excavation analysis and facilitating contact with Scottish Power.

## Data Availability Statement

Data openly available in a public repository that issues datasets with DOIs

