## Supplementary Material for "Early Neolithic forest farming at Seven Springs, Martlesham, Suffolk inferred from sedaDNA and pollen"

### Text S1- Extended Methodology

#### Sampling & Sedimentology

Sampling was performed in July 2017, using six 50ml tubes, with the lids and threads sterilised with bleach, that were inserted into the sediment. Alongside, a 70cm u-channel was taken for pollen and sedimentological analyses. Loss on Ignition (LOI) analysis was conducted following the standard methodology (Heiri *et al.* 2001) and consisted of 22 samples of 2mg sediment, subsampled at 1cm resolution from the u-channel. Samples were heated in an oven for 12 hours at 50°C to estimate moisture content, burned at 550°C for two hours to estimate organic content and finally re-ignited at 950°C for four hours to estimate carbonate content. Magnetic susceptibility analysis was undertaken using a Bartington MS2 meter at 1cm resolution on the u-channel using the MS2B Dual Frequency Sensor for HF and LF, and the MS2K downcore Surface Sensor. To support the sedimentology pXRF was performed using a NITON XL3 analyzer at 1cm resolution on the u-channel, both the geochemical and soil settings were used to obtain a final suite of 44 elements.

#### ***Seda*DNA**

*Metabarcoding*

*Sed*aDNA extraction followed protocols documented by Hudson *et al.* (2022) using the QIAGEN DNeasy Powersoil Kit and was performed in a dedicated ancient DNA laboratory at the University of Southampton. Metabarcoding PCR amplification targeted plants, using primers for the P6 loop of *trn*L UAA intro of the chloroplast genome (Taberlet *et al*. 2007), and mammals using primers for the mitochondrial 12S (batra) and 16S (mamP007) rRNA regions (Giguet-Covex *et al.* 2014; Valentini *et al.* 2016). For 12S however, we used a modified reverse primer with sequence GTAYRCTTACCWTGTTACGACTT, and used a dual-blocking approach, which consisted of the batra_blk forward blocker and a reverse blocker with sequence GTTACGACTTGTCTCCTCTATATAAATGCGAT-C3. PCR recipes and reaction conditions followed Voldstad *et al.* (2020) for plants and Garces-Pastor et al. (2022) for mammals. Four negative extraction controls, two PCR negative controls and one positive control were carried out. Eight individually tagged PCR replicates were made for each sample to increase the chance of detecting taxa represented by low quantities of sedaDNA, as well as to increase confidence in taxa being detected. Paired-end sequencing was performed on an Illumina NextSeq-550 at the Genomics Support Centre Tromsø (GSCT) at UiT- The Arctic University of Norway. All next-generation sequence data were aligned, filtered and trimmed using the OBITools software package (Boyer *et al.* 2016) using similar criteria as Alsos *et al.* (2020). Resulting barcodes were assigned to taxa using the *ecotag* program and, for plants, using four independent reference datasets. One reference contained arctic (Sønstebø *et al*. 2010) and boreal (Willerslev *et al*. 2014) vascular plants as well as bryophytes from the circumpolar region (Soininen *et al*. 2015) (ArcBorBryo, n=2280 sequences of which 1053 are unique), one the NCBI nucleotide database (January 2021 release), one the PhyloAlps database (Garces-Pastor *et al.* 2022) and finally the PhyloNorway component of the NorBOL database (Alsos *et al.*2022). For mammals, only the NCBI nucleotide database was used.

The resulting identifications were merged and filtered, for plants, retaining barcode sequences if they were identified to >98% in at least two reference sets and had at least 10 reads across the entire dataset, and for mammals if they were identified to 100%. False positives relating to common PCR errors and food contaminants were removed based on 'blacklists' built up from previous research at The Arctic University Museum of Norway, as well as taxa identified above family level. Manual checks were also completed for a few taxa that were not ecologically plausible. For the last step of filtering, the frequency of PCR replicates in samples compared to negative controls was examined. Sequences were retained if they had an overall frequency of PCR replicates in samples at least twice as high as in that in negative controls. We present the proportions of PCR replicates 1-8 and total reads. Where a reference dataset was unable to assign to specific species due to sequence sharing, we have listed all species that share the sequence.

*Shotgun Metagenomics*

Aliquots of the same *seda*DNA extracts used for metabarcoding were sent to the paleogenetics laboratories at AWI Potsdam (Germany) for shotgun metagenomics sequencing. Three samples -M90-56cm, M91-52cm, M92-54cm, and a library blank control were prepared with the single-stranded library method specified for degraded ancient DNA (Gansauge et al. 2017, Gansauge & Meyer 2013) with modifications described in Schulte et al. (2020). DNA extracts were quantified with Qubit 4.0 fluorometer (Invitrogen) and 30 ng of DNA were used as the starting material. After adapter ligation, an indexing PCR with 10 to 13 PCR cycles and unique combinations of P5 and P7 Illumina primers was performed. Then, PCR products were purified with the MinElute PCR Purification Kit (Qiagen, Germany), DNA concentration was measured with Qubit 4.0 fluorometer (Invitrogen) and libraries fragment sizes were checked on the Agilent TapeStation using the D1000 ScreenTape (Agilent Technologies USA). Finally, sample libraries were pooled equimolarly to 20 nM, while 1 µl of library blank was added to the pool. Pooled samples were sequenced with a customized sequencing primer CL72 on a NextSeq2000 Illumina device using paired-end mode (2x100 bp) at AWI Bremerhaven (Germany). After sequencing, overlapping paired-end reads were merged and adapters trimmed using SeqPrep2 (https://github.com/jeizenga/SeqPrep2), while using a minimum overlap of 15 nucleotides and only retaining reads that merged and were ≥35 bp. Low complexity reads were then removed with a DUST score of ≤1 using SGA preprocess (Simpson *et al.* 2012). For the taxonomic classification of reads, we used the k-mer-based classifier Kraken2 v. 2.0.8-beta86 in (confidence threshold of 0.8). We downloaded the National Center for Biotechnology Information (NCBI) non-redundant nucleotide database (ftp://ftp.ncbi.nlm.nih.gov/blast/db/FASTA/nt.gz; downloaded in June 2021) and the NCBI taxonomy (obtained using kraken2-build in June 2021) for taxonomic classification and built the Kraken2 database. This classification of the shotgun data was used for comparison between the shotgun and metabarcoding datasets. The Kraken2 analysis suggested the presence of *Corylus*, *Betula*, *Bos*, and *Ovis* within the samples, which were chosen for additional validation analyses.

*Ancient DNA Damage Analysis, and* Bos *Mitogenomic Comparison*

The pre-processed data was aligned against two composite references consisting of 3-4 reference genomes: (1) Mammal: sheep, cow, wolf, human, (2) Plant: white birch, hazel, cushion willow (Table S1). To each composite reference, we added a list of conserved sequences (v1; https://github.com/pheintzman/metagenomics), which together with wolf/human or willow acted as decoy references for competitive mapping purposes (Feuerborn *et al.* 2020). Reads were independently mapped against the two composite references using bwa v0.7.17 (Li and Durbin 2010) with the seed disabled (-l 1024) (Schubert *et al.* 2012). Reads that had a mapping quality score of <25 were removed using samtools v1.9 (Li 2009). Duplicated reads were removed using dedup v0.12.7 (Peltzer *et al.* 2016). Relative read mapping frequencies were calculated against each component reference genome (in reads mapped per Mb). Mean fragment lengths were calculated using samtools and R v3.6. The ancient DNA damage profiles were visualised using mapDamage v2.2.0 (Jonsson *et al.* 2013) with the –single-stranded and –merge-reference-sequences. The mean damage rate was calculated by subtracting the mean C-to-T transition rate at 5' bases at positions 11-25 from the C-to-T transition rate at the 5' first position. To test whether DNA damage rates were statistically robust, we used pydamage v0.80 (Borry et al. 2021), which also considers damage rates beyond first position. In pydamage, we simultaneously considered all contigs/scaffolds from the same genome in calculations (--group flag).

To test evenness of coverage between scaffolds within each component reference genome, we calculated the $R^2$ fit between count of reads aligned and scaffold length for each component reference genome and sample combination, using data calculated by samtools idxstats. As a greater amount of data aligned to reference genomes within the plant composite reference, we used scaffolds ≥1 Mb, whereas we used ≥10 Mb for the mammal composite reference. The tests of evenness of coverage highlighted that select scaffolds within the mammal composite reference had greater counts of reads aligned than expected in both samples (e.g. NC_037341 in the cow genome, CM029821 in sheep). Inspection of the BAM files showed reads concentrated in small regions and that could be identified as likely originating from plants based on NCBI blastn nucleotide queries. We therefore sought to remove any reads mapped to the mammal composite reference that may have had a non-mammalian origin. For this, we extracted reads aligned to the mammal composite reference using PicardTools v2.21.1 SamToFastq (http://broadinstitute.github.io/picard) and the FASTX toolkit v0.0.14 (http://hannonlab.cshl.edu/fastx_toolkit). We aligned the reads to the NCBI nucleotide using NCBI blast+ v2.12.0 blastn (-task blastn) (Camacho *et al.* 2008) and retained any that had a hit to non-Eutherian taxids (using -negative_taxids). The taxid list, consisting of all Eutherian taxids, was generated using TaxonKit v0.12.0 list (Shen & Ren 2021) with the NCBI taxdump file downloaded on 2022-06-21 and the taxid set to 9347 (Eutheria). We then removed any reads that had a blast hit from the mammal composite reference BAM files and regenerated relative read mapping frequencies, mean fragment lengths, ancient DNA damage profiles and rates, and evenness of coverage statistics.

Lastly, to test whether our cow data likely represented domestic cow or wild aurochs, we leveraged recent findings that show Holocene aurochsen from west Europe carry mitochondrial haplogroup P (Achilli et al. 2009; Rossi et al. 2024), which is diverged from haplogroups found in taurine cattle. The latter has haplotypes related to some Asian aurochs, consistent with the domestication history of cattle. On the other hand, other Asian aurochs carry haplogroups that are outgroup to those found in both taurine cattle and west European aurochs (Rossi et al. 2024). We first extracted metagenomics reads that aligned to the Cow mitogenome (NC_006853) within our mammal composite reference, as well as the 16S and 12S Bos sequences derived from metabarcoding. We compared these reads to a panel of 20 *Bos* mitogenomic sequences: the Bovine Reference Sequence (*B. taurus*; V00654), which is very similar to haplogroup T3 and typical of West European taurine cattle, and the 19 available mitochondrial genomes for aurochs (*B. primigenius*) from NCBI Genbank, including those from Europe (n=8), Asia (n=10), and one of unclear provenance (Table S2).

**Pollen**

Standard pollen extraction techniques (Moore and Webb 1978; Moore *et al.* 1992) were used on samples of 1.5ml volume taken from the monolith profile at 2cm intervals. Where possible, a sum of *c.* 300 pollen grains per sample was identified and counted using an Olympus biological research microscope. Marsh taxa and fern spores were counted outside of the basic pollen sum. A pollen diagram (figure **) was plotted using Tilia with percentages calculated as follows:

Sum = % total dry land pollen but including *Alnus* and *Salix*

Marsh/aquatic herbs = % tdlp + sum of marsh/aquatics
Spores = % tdlp + sum of spores
Misc. = % tdlp + sum of misc. taxa.

Taxonomy, in general, follows that of Moore and Webb (1978) modified according to Bennett *et al.* (1994) for pollen types and Stace (1991). A substantial comparative collection of British and European pollen types was available to assist any identification problems. These procedures were carried out in the Palaeoecology Laboratory of the School of Geography and Environment, University of Southampton.

### Text S2- Extended Sedimentological and Environmental Results

#### LOI, pXRF and Mag Sus

From the LOI, it can be seen that water content increases through the marl, remains high throughout the peaty soil within which the wood remains are found, then dips slightly into the upper minerogenic peat (Figure S1). Water content closely mirrored organic content, and the upper marl horizons were shown to gradually increase in organics. Intra-horizon elemental variation was low, but a subtle iron pan was detected at the top of the marl.

#### SedaDNA Metabarcoding and Pollen

The 6 samples amplified for the trnL P6 loop of the plant chloroplast genome produced 2,787,181 reads which after post-identification filtering was reduced to 2,150,865 reads with 87 taxa, 13 to family, 43 to genus and 31 to species (Dataset S1). Target DNA preservation was good, with an average of approximately 77% of reads retained after all pre and post-identification filtering (Text S1). Overall percentages of reads were dominated by woodland taxa at every depth but the forb assemblage was large and diverse, further discussion of the sedaDNA can be found in Text S2.

From the combined mitochondrial 12S and 16S rRNA PCR amplifications of the same samples (Dataset S2), 386,887 reads were obtained. The majority of sequences were those of prokaryotes and after the removal of these alongside any *Homo* DNA, 10,023 reads originating from *Bos, Ovis, Capra* and *Sus* were identified. After comparison with *Bos primigenius* sequences, the *Bos* 12S and 16S sequences were deemed to be representative of domestic cattle and not aurochsen, as they matched haplotype T3B, which is found in modern domestic breeds that are derived from Anatolian aurochsen. The 12S primers were more successful, amplifying all four of the mammal taxa, whilst the 16S primers amplified only *Bos* and *Ovis*. *Ovis* and *Capra* are genetically distinct, differing by 12 substitutions in the 12S target sequence.

One feature that stands out is the breadth of forb taxa represented (>60), most of which are seen across different samples in high numbers of PCR replicates which increases confidence in their presence (Chen and Ficetola 2020). However, the limited number of reads of almost all herb types (Figure 5), with the exception of wetland herbs, suggesting most were not likely abundant on-site but marginal or upstream. Whilst plant *seda*DNA deposition is imperfectly understood (Fonseca 2018), it usually represents predominantly locally growing taxa (Alsos *et al.* 2018), and one reason for the high numbers of taxa could be the sedimental context at Martlesham being that of a palaeochannel with active spring activity. The *sed*aDNA read data is heavily dominated by alder, reflecting the localised bias to species within the palaeochannel, and without the addition of the pollen a clearer picture of the nature of the surrounding deciduous woodland would not be possible. Conversely, the high pollen counts from the alder, lime, oak and hazel woodland may ‘swamp’ signatures of the smaller populations of herbs seen in the *sed*aDNA and it is this reciprocity that makes pollen and *seda*DNA complimentary techniques.

The birch and pine seen in low counts in the pollen assemblage (<5%) were not growing near to the sample site as they are not seen in the *seda*DNA and are therefore likely to be predominantly windblown. Another significant different between the two assemblages is the extra-local presence of maple (*Acer*) in upper samples of the *seda*DNA, a taxon often unrepresented in pollen counts but one identified in the site preserved wood assemblage and so definitely present on site.

**SedaDNA Shotgun Metagenomics and Ancient DNA Damage Analysis**

**Evidence for the presence of hazel (*Corylus*)**

In both samples, we observed a 2.0 to 3.3-fold increase in the proportion of reads mapping to the hazel genome as compared to the birch genome (after controlling for relative genome sizes; Table S3). In the negative control, three reads aligned to the birch genome as compared to 52,000-109,000 in the samples, whereas no reads aligned to either of the other two plant genomes. The $R^2$ fit between counts of reads mapped and scaffold length was higher for hazelnut (0.93-0.94) than for birch (0.79-0.83), consistent with hazel providing a more appropriate reference genome for the data than birch (Table S4). Reads mapped to all three plant genomes showed an excess of C-to-T transitions at the 5’ first position as compared to the average at positions 11-25, which is statistically significant (all with a predicted accuracy of 1 and $p<0.001$; Table S5; Figure S2-S4). This is consistent with the presence of hazel, birch, and willow ancient DNA in both sediment samples. However, the rate of C-to-T transitions, as well as other substitution types, across positions 11-25 is greater for reads aligned to birch (0.027-0.028) than to hazel (0.015-0.018) (Table S5; Figure S2-S3). Altogether, the analyses show that there is a population of reads that is more closely related to the hazel genome than to birch.

**Evidence for sheep (*Ovis*) and cow (*Bos*) presence**

We mapped 79-120 reads to the sheep genome and 84-163 to the cow genome. After duplicate removal and BLAST filtering, we retained 38-44 and 36-67 reads to the sheep and cow genomes, respectively. No reads from the negative control aligned to either target genome (Table S6). In both sediment samples, the retained reads for both sheep and cow showed an excess of C-to-T transitions at the 5' first position as compared to the average at positions 11-25, which is statistically significant (all with a predicted accuracy of 1 and p=0.002-0-003; Table S7). This suggests a signal of ancient DNA damage for both taxa in these samples. In contrast, reads mapped to the decoy genomes in samples, or to all genomes in the negative controls, show no evidence of ancient DNA damage (p=0.081-1.000; Table S7; Figure S5), consistent with a modern origin for these reads. Visual inspection of the ancient DNA damage plots suggests a higher rate of mismatch for reads aligned to the sheep genome in JK135L-3, as compared to JK135L-2 and reads aligned to the cow genome for both samples (Figure S6-S7). After applying the BLAST filter, we observed an increase in the $R^2$ fit between counts of reads mapped and scaffold length, consistent with broadly correct read assignment (Table S8). The exception was JK135L-3 reads aligned to the sheep genome, whereby the $R^2$ fit decreased after BLAST filtering. Overall, the $R^2$ values for sheep and cow were low, as compared to plants, which likely results from their far smaller data set sizes.

**Bovine mitogenomic comparison**

We extracted nine metagenomics reads and two metabarcodes that aligned to the cow mitochondrial genome across both samples (Table S9). Six reads were either identical to the corresponding regions in the *Bos* mitogenomic sequence panel (n=4) or only carried a proximal C-to-T/G-to-A allele not present in the panel and likely deriving from ancient DNA damage (n=2). The remaining two reads and two metabarcodes carried eight alleles that matched either the Bovine Reference Sequence, west European aurochs, Asian aurochs, or were found at ~50% frequency within one of the aurochs groups. Seven of the alleles matched the Bovine Reference Sequence, whereas the European and Asian aurochs were both supported by five alleles. The single allele not found in the Bovine Reference Sequence is shared by all aurochs and likely reflects an ancestral state, as other taurine diagnostic alleles were present in the same metabarcode sequence (Table S9). This analysis is therefore consistent with the *Bos* sedaDNA data reads originating from taurine cattle and not aurochs.

### Text S3- Wood

The information here is taken from the preliminary assessments as full analysis of all the site material is pending post-excavation analysis.

**Preserved Wood from Palaeochannel 1**

**Context 231569 (peat at aurochs core location).**

Eight pieces of wood and timber. Five fragments of *Alnus* spp. root and three bark fragments not identifiable to species. All pieces have a high degree of fragmentation, though largely refittable. No evidence for woodworking or modification.

**Context 231799**

1123 pieces of wood and timber. Of these, 549 are fragments of bark, nine of which had wood present and could be identified to species (eight *Alnus* spp. one *Quercus* spp.). There are 508 fragments of root, all of which are *Alnus* spp. except for one that was too degraded for identification. Five of these were root stumps with multiple side shoots originally present. There are 30 pieces of branch wood of which seventeen are *Quercus* spp. and thirteen *Alnus* spp. 11 fragments of small diameter roundwood from smaller branches or twigs were recorded, all of which are *Alnus* spp. 9 sections or fragments of roundwood log are present, of which six are *Alnus* spp., one *Quercus* spp. and two were too degraded for identification. None of the above had any evidence for woodworking.

Just 15 pieces of wood had potential evidence for human modification and in some cases, these are possibly, but not definitely worked. Six of these are probably offcuts, based on having conversion and cross-sectional shapes that would be difficult to recreate by natural processes of decay or damage. Three are tangentially faced *Alnus* spp. and three are radially and tangentially faced *Quercus* spp. fragments. There are three possible fragments of board, two of which are tangentially faced *Alnus* spp., one radially faced *Quercus spp.* Five possible post or stake tips are present of which three are *Quercus* spp., one is *Fraxinus excelsior L.* and one *Ulmus* spp has possible stone axe hewing marks. Finally, there is a single peg or small stake cut from *Quercus* spp. heartwood.

**Preserved Wood from Comparable Contexts**

**Context 231814 (peat).**

A total of 117 pieces of wood and timber were recovered. Of these, 34 are fragments of bark, three of which had wood present and could be identified to species (two *Alnus* spp., one

*Quercus* spp.). There are 65 fragments of root, all of which are *Alnus spp.* Two of these were root stumps with multiple side shoots originally present. There are 8 pieces of branch wood of which five are *Alnus* spp. and three *Quercus* spp. 5 sections or fragments of roundwood log are present, of which one is *Alnus* spp., three *Quercus* spp. And one *Fraxinus excelsior L.* None of the above had any evidence for woodworking. There are also 2 bags labelled as timbers or wood fragments but which on examination and cleaning proved to be compressed blocks of peat with no actual wood present.

**Context 231826.**

A total of 14 pieces of wood and timber was recovered from this context. Of these, 5 are fragments of bark. There are 6 fragments of root, all of which are *Alnus* spp. There are 3 pieces of *Alnus* spp roundwood. None had any evidence of woodworking.

### Text S4 Animal Bones

The information here is taken from the preliminary and amended assessments as full analysis of all the site material is pending post-excavation analysis.

**Observations on species present:**

The assemblage from Palaeochannel 1 contains bones from bovids, equids, sheep/goat, red deer, roe deer, alongside the remains of an aurochs (*Bos primigenus*).

*Bovids (Cattle/Auroch):* These dominated the overall site assemblage with numerous elements of cattle bones throughout, with a range of elements and ages noted. Several pathologies were noted which should provide information on health and husbandry; some working cattle are indicated. From Palaeochannel 1, three bones were identified from the lower peats and part of a skull and a pair of horncores from an auroch were recovered from peat (231569) in Palaeochannel 1 with a head and horncore width of approximately 1m; the size of the horncores indicating a bull. There is a possible cut mark on the skull, suggesting it may have at least been skinned. Other bovid remains in the assemblage are of a large size and may also be from aurochs.

*Equid (Horse): is present* in Palaeochannel 1. Butchering was noted on some equid remains, showing some use of this species for skin and meat. Several pathologies were seen, which includes probable bit-wear and hock-arthritis.

*Sheep/goat:* In the whole site assemblage, most sheep/goat bones are sheep, with goat present in small numbers. From Palaeochannel 1, only one sheep/goat bone is identified in the upper peat fills.

*Deer:* One red deer mandible was identified from the lower peats. Two Roe Deer antlers from the basal sands/marl (233427), considered in the field to be parts of possible antler headdresses, were selected for an enhanced assessment. The two antlers from context (233427 - sand) were found separately towards the close of the excavation. Roe Deer antler one is a complete right side antler with part of the frontal bone of the skull attached. The lowest tine has lost the tip, which is most likely to have occurred when the deer was in a battle during the rut. There is a hole in the frontal bone of this piece, but this is a natural blood vessel hole and not a human modification. There are numerous clear, often crossing, cut marks on the frontal bone. The butchering and polishing of the Roe Deer antlers from Palaeochannel 1 strongly suggest that these items were selected and prepared for use. The evident polishing on all three antlers further suggesting that they were regularly handled, rather than used for decorative purposes.

**Table S1- Reference Genomes for Shotgun Sequencing genomic comparisons.**

| | | Species | | Genome assembly | | | | |
|---|---|---|---|---|---|---|---|---|
| **Composite** | **Purpose** | **Common name** | **Latin name** | **Name** | **NCBI accession** | **Level** | **Size (bp)** | **Notes** |
| Mammal | Target | Sheep | *Ovis aries* | CAU_O.aries_1.0 | GCA_017524585.1 | chromosome | 2,653,843,355 | |
| Mammal | Target | Cow | *Bos taurus* | ARS-UCD1.3 / bosTau9 | GCF_002263795.2 | chromosome | 2,711,209,831 | |
| Mammal | Decoy | Wolf | *Canis lupus* | mCanLor1.2 | GCA_905319855.2 | chromosome | 2,447,463,909 | |
| Mammal | Decoy | Human | *Homo sapiens* | GRCh37 / hg19 | GCA_000001405.1 | chromosome | 3,137,161,264 | Includes mtDNA genome (16571 bp) |
| Plant | Target | White birch | *Betula pendula* | Bpev01 | GCA_900184695.1 | scaffold | 435,914,794 | |
| Plant | Target | Hazel | *Corylus avellana* | CavTom2PMs-1.0 | GCA_901000735.2 | chromosome | 369,779,143 | |
| Plant | Decoy | Cushion willow | *Salix brachista* | ASM907833v1 | GCA_009078335.1 | chromosome | 339,587,529 | |

**Table S2- Aurochs (*B. primigenius*) mitogenomes used for determining the taxonomic origin of *Bos* reads.**

| **Genbank accession** | **Country** | **For_analysis** |
|---|---|---|
| GU985279 | UK | European aurochs |
| JQ437479 | Poland | European aurochs |
| MF169213 | Scandinavia | European aurochs |
| MF169212 | Scandinavia | European aurochs |
| MF169211 | Scandinavia | European aurochs |
| MW689251 | Spain | European aurochs |
| MW689250 | Spain | European aurochs |
| MW689249 | Spain | European aurochs |
| OQ160855 | China | Asian aurochs |
| OQ160854 | China | Asian aurochs |
| OQ160850 | China | Asian aurochs |
| OQ160849 | China | Asian aurochs |
| OQ160848 | China | Asian aurochs |
| OQ160846 | China | Asian aurochs |
| OQ160845 | China | Asian aurochs |
| OQ160844 | China | Asian aurochs |
| OQ160843 | China | Asian aurochs |
| OQ160842 | China | Asian aurochs |
| OL960715 | Unknown | Not used |

**Table S3- Comparative Mapping to Reference Genomes**

| | Total | Birch | Hazel | Willow | conserved_seqs |
|---|---|---|---|---|---|
| Genome size | 1,145,281,963 | 435,914,794 | 369,779,143 | 339,587,529 | 497 |
| | Reads mapped per Mb across entire ref. genome | | | | |
| Sample | Total | Birch | Hazel | Willow | conserved_seqs |
| JK135L-2 (52cm) | 104.07 | 99.55 | 199.03 | 6.45 | |
| JK135L-3 (54cm) | 306.35 | 207.52 | 684.04 | 21.95 | |
| JK135L-4 (Blank) | 0.00 | 0.01 | | | |

**Table S4- $R^2$ fit between count of reads aligned and scaffold length for each component reference genome and sample combination**

| | Scaffold length vs. reads aligned ($R^2$; across scaffolds >1 Mb) | | | | |
|---|---|---|---|---|---|
| **Sample** | **Total** | **Birch** | **Hazel** | **Willow** | **conserved_seqs** |
| JK135L-2 (52cm) | na | 0.7974 | 0.9318 | 0.4715 | |
| JK135L-3 (54cm) | na | 0.8344 | 0.9392 | 0.4219 | |
| JK135L-4 (Blank) | na | | | | |

**Table S5- Plant ancient DNA damage analyses.** From top to bottom: (top) mean C-to-T transition rate at the 5’ first position for plants after correction for (upper middle) the mean C-to-T transition rate at positions 11-25. Results from pydamage are shown as (lower middle) q-value, with a statistically significant result of the presence of aDNA damage being q<0.05, and (bottom) the pydamage estimated model accuracy.

| | **Plants: corrected aDNA damage rate (C -> T, 5' base 1)** | | | |
|---|---|---|---|---|
| Sample | **Birch** | **Hazel** | **Willow** | **conserved_seqs** |
| JK135L-2 (52cm) | 0.1205 | 0.1673 | 0.1590 | |
| JK135L-3 (54cm) | 0.1527 | 0.1884 | 0.1339 | |
| JK135L-4 (Blank) | 0.0000 | | | |

| | **Plants: mean divergence (C -> T, 5' bases 11-25)** | | | |
|---|---|---|---|---|
| Sample | **Birch** | **Hazel** | **Willow** | **conserved_seqs** |
| JK135L-2 (52cm) | 0.0274 | 0.0176 | 0.0206 | |
| JK135L-3 (54cm) | 0.0283 | 0.0147 | 0.0183 | |
| JK135L-4 (Blank) | 0.0000 | 0.0000 | 0.0000 | |

| | **Plants: pydamage aDNA damage rate (q-value)** | | | |
|---|---|---|---|---|
| Sample | **Birch** | **Hazel** | **Willow** | **conserved_seqs** |
| JK135L-2 (52cm) | <0.001 | <0.001 | <0.001 | |
| JK135L-3 (54cm) | <0.001 | <0.001 | <0.001 | |
| JK135L-4 (Blank) | 1.0000 | | | |

| | **Plants: pydamage aDNA damage rate (model accuracy)** | | | |
|---|---|---|---|---|
| Sample | **Birch** | **Hazel** | **Willow** | **conserved_seqs** |
| JK135L-2 (52cm) | 1.0000 | 1.0000 | 1.0000 | |
| JK135L-3 (54cm) | 1.0000 | 1.0000 | 1.0000 | |
| JK135L-4 (Blank) | 0.9940 | | | |

**Table S6- Reads Mapping to Animal Reference Genomes after Duplicate Removal and BLAST Filtering**

| | Animals: non-Eutherian reads excluded | | | | | |
|---|---|---|---|---|---|---|
| Sample | **Total** | **Sheep** | **Cow** | **Wolf** | **Human** | **conserved_seqs** |
| JK135L-2 (52cm) | 3300 | 38 | 36 | 1 | 3225 | 0 |
| JK135L-3 (54cm) | 9222 | 44 | 67 | 16 | 9095 | 0 |
| JK135L-4 (Blank) | 215 | 0 | 0 | 2 | 213 | 0 |

**Table S7- Animal ancient DNA damage analyses.** From top to bottom: (top) mean C-to-T transition rate at the 5' first position for plants after correction for (upper middle) the mean C-to-T transition rate at positions 11-25. Results from pydamage are shown as (lower middle) q-value, with a statistically significant result of the presence of aDNA damage being q<0.05, and (bottom) the pydamage estimated model accuracy.

| | **Animals: corrected aDNA damage rate (C -> T, 5' base 1)** | | | | |
|---|---|---|---|---|---|
| Sample | **Sheep** | **Cow** | **Wolf** | **Human** | **conserved_seqs** |
| JK135L-2 (52cm) | 0.2500 | 0.3667 | 0.0000 | 0.0127 | 0.0000 |
| JK135L-3 (54cm) | 0.3222 | 0.1622 | 0.0000 | 0.0031 | 0.0000 |
| JK135L-4 (Blank) | | | 0.0000 | -0.0041 | |
| | **Animals: mean divergence (C -> T, 5' bases 11-25)** | | | | |
| Sample | **Sheep** | **Cow** | **Wolf** | **Human** | **conserved_seqs** |
| JK135L-2 (52cm) | 0.0000 | 0.0083 | 0.0000 | 0.0054 | |
| JK135L-3 (54cm) | 0.0111 | 0.0044 | 0.0000 | 0.0018 | |
| JK135L-4 (Blank) | 0.0000 | 0.0000 | 0.0000 | 0.0041 | |

| | **Animals: pydamage aDNA damage rate (q-value)** | | | | |
|---|---|---|---|---|---|
| Sample | **Sheep** | **Cow** | **Wolf** | **Human** | **conserved_seqs** |
| JK135L-2 (52cm) | 0.0020 | 0.0030 | 1.0000 | 0.0900 | |
| JK135L-3 (54cm) | 0.0020 | 0.0030 | 1.0000 | 0.0810 | |
| JK135L-4 (Blank) | | | 1.0000 | 0.9980 | |
| | **Animals: pydamage aDNA damage rate (model accuracy)** | | | | |
| Sample | **Sheep** | **Cow** | **Wolf** | **Human** | **conserved_seqs** |
| JK135L-2 (52cm) | 1.0000 | 1.0000 | 1.0000 | 1.0000 | |
| JK135L-3 (54cm) | 1.0000 | 1.0000 | 1.0000 | 1.0000 | |
| JK135L-4 (Blank) | | | | 1.0000 | |

### Table S8- Reads mapped and scaffold length for animals with BLAST filtering applied

| | Scaffold length vs. reads aligned ($R^2$; across scaffolds >10 Mb): WITH BLAST FILTERING | | | | | |
|---|---|---|---|---|---|---|
| **Sample** | **Total** | **Sheep** | **Cow** | **Wolf** | **Human** | **conserved_seqs** |
| JK135L-2 (52cm) | na | 0.4973 | 0.1725 | | 0.8019 | |
| JK135L-3 (54cm) | na | 0.0608 | 0.2099 | 0.1028 | 0.9224 | |
| JK135L-4 (Blank) | na | | | | 0.5173 | |

**Table S9- Bovine mitogenomic comparison- reads that aligned to the *Bos* mitochondrial genome.** Coordinate positions are derived from the Bovine Reference Sequence.

| Data type | Sample | Sequence | Length (bp) | Start position | MQ score | Retained after BLAST filtering? | Allelic summary | Allele in seda DNA | SNP position | Allele in taurine cattle (BRS) | Allele in aurochs (west Europe) | Allele in aurochs (Asia) | aDNA damage? | Note |
|---|---|---|---|---|---|---|---|---|---|---|---|---|---|---|
| Metagenomic read | JK135L-2 | AATTGGCCTAACAACAAATA TACTAACAATATACCAATGA TGACGAGATGTTATCCGAG AAAGCACCTTCCAAGGGCA | 78 | 9101 | 37 | no | none | na | na | na | na | na | no | |
| Metagenomic read | JK135L-2 | AAAGTATGCAAGAACTGCT AATTCTATGCTCCCATATCT AATAGTATGGCT | 51 | 11980 | 37 | no | Taurine / Asia auroch | T | 12016 | T | C | T | no | |
| Metagenomic read | JK135L-2 | CATCTTAGCCCTAGAAATCA GTAATATAACTAAAAATCTA AAATATCACTACCCCTCAA | 59 | 13602 | 37 | no | Taurine / Euro auroch | C | 13620 | C | C | T | no | |
| Metagenomic read | JK135L-3 | AGATAATTACATAAACAAAA TTATTCGCCAGAGTACTACT AGCAACAGCTTAAAACTCA AAGGACTT | 67 | 884 | 37 | no | none | na | na | na | na | na | no | |
| Metagenomic read | JK135L-3 | TACCTGCCCAGTGACAACT GTTTAACGGCCGCGGTATC CTGACCGTGCAAAGG | 53 | 2336 | 37 | no | none | na | na | na | na | na | yes, second position | |
| Metagenomic read | JK135L-3 | CCTAGGAGTCCTATTTATAC TAGCCATATCAAGCCTAGC CGTATACTCCATTT | 53 | 3397 | 37 | no | none | na | na | na | na | na | yes, first position | |
| Metagenomic read | JK135L-3 | TTTTATCATCTTTCAACTAA AAGTTTCAAAACACAACTTT TATCACAATCCAGAACT | 57 | 8188 | 37 | yes | Taurine / Euro auroch | T | 8188 | T | T | C | no | First position, so could be aDNA damage |
| | | | | | | | Taurine | T | 8236 | T | C | T | no | |
| Metagenomic read | JK135L-3 | GTTATAGCAGCCCTAACAA TCCTCAACTCACATTTTACA TTAGCTAGCA | 49 | 10362 | 37 | yes | none | na | na | na | na | na | no | |
| Metagenomic read | JK135L-3 | ATACCCATTCATTATACTCT CCCTATGAGGCATAATTATA ACCAGCTCAATCTG | 54 | 11302 | 37 | no | none | na | na | na | na | na | no | |
| Metabarcoding amplicon | Both (12S) | CCTCAAATAGATTCAGTGC ATCTAACCCTATTTAAACGC | 60 | 1281 | *na* | *na* | not diagnostic | C | 1307 | C | C | C / T | no | |

| | | | | | | | | | | | | | |
|---|---|---|---|---|---|---|---|---|---|---|---|---|---|
| | | ACTAGCTACATGAGAGGAG AC | | | | | | | | | | | |
| Metabarcodin g amplicon | Both (16S) | TTAACTAACCAACCCAAAG AGAATAGATTTAACCATTAA GGAATAACAACAATCTCCA TGAGTTGGTAGTTTC | 73 | 2533 | *na* | *na* | not diagnostic | T | 2585 | T | C / T | T | no |
| | | | | | | | Taurine | G | 2558 | G | A | A | no |
| | | | | | | | Euro / Asia auroch | A | 2536 | C | A | A | no |

**Table S10- Radiocarbon Date Table: Calibrated on OxCal 4.4 using IntCal21 (Reimer *et al.* 2020).**

| **Lab ID** | **Context [Location]** | **Context description** | **Material dated** | **$\delta^{13}C$ (‰)** | **Radiocarbon age (BP)** | **Calibrated date (68%)** | **Calibrated date (95%)** |
|---|---|---|---|---|---|---|---|
| SUERC-79765 | 231569 | Peat infill of Palaeochannel 1 | Animal bone: Auroch | -23.4 | 5502 ± 32 | 4440-4330 cal. BC | 4450-4260 cal. BC |
| SUERC-81020 | 231799 | Peat infill of Palaeochannel 1 with preserved wood remains | Waterlogged wood: Alnus spp. | -28.7 | 4699 ± 27 | 3520-3380 cal. BC | 3620-3370 cal. BC |
| SUERC-81019 | 231799 | Peat infill of Palaeochannel 1 with preserved wood remains | Waterlogged wood: *Alnus spp.* | -29.3 | 4471 ± 27 | 3330-3090 cal. BC | 3340-3030 cal. BC |
| SUERC-81021 | 231814 | Peat infill of Palaeochannel 1 | Waterlogged wood: *Fraxinus excelsior* | -30.3 | 4358 ± 27 | 3010-2920 cal. BC | 3080-2900 cal. BC |
| SUERC-81024 | 233367 | Peat infill of Palaeochannel 1 | Waterlogged wood: *Alnus spp.* | -27.7 | 1634 ± 27 | cal. AD 410-530 | cal. AD 380-540 |

**Figure S1- pXRF and LOI data**

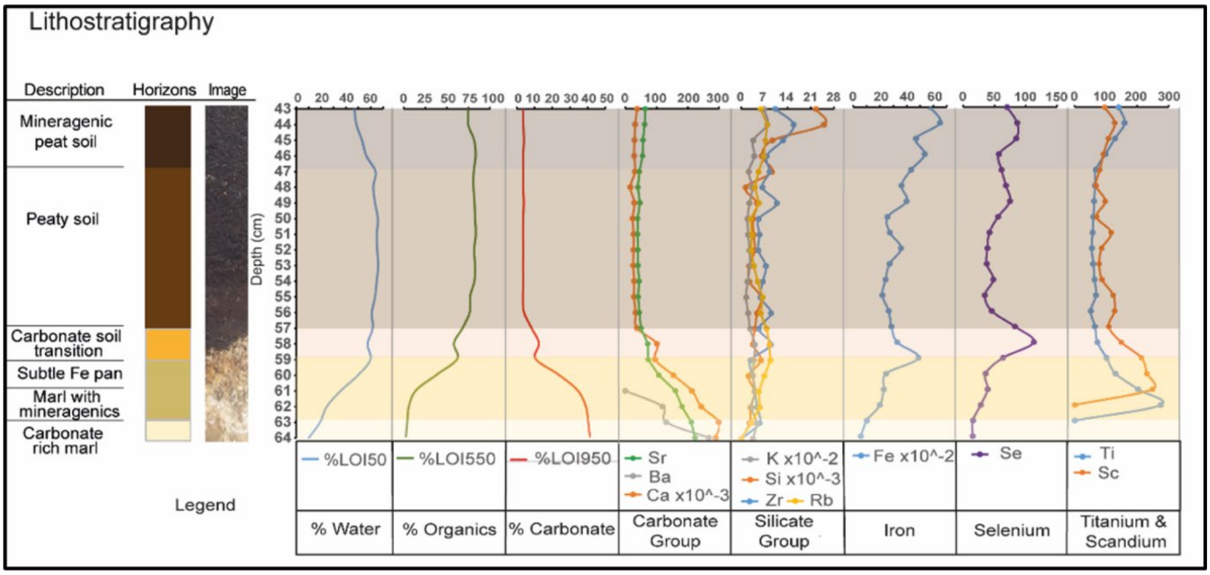

Figure S1 Lithostratigraphic data from the sampled u-channel from Palaeochannel 1

**Figure S2- Read length distributions and ancient DNA deamination plots for reads mapped to the *Betula pendula* genome**

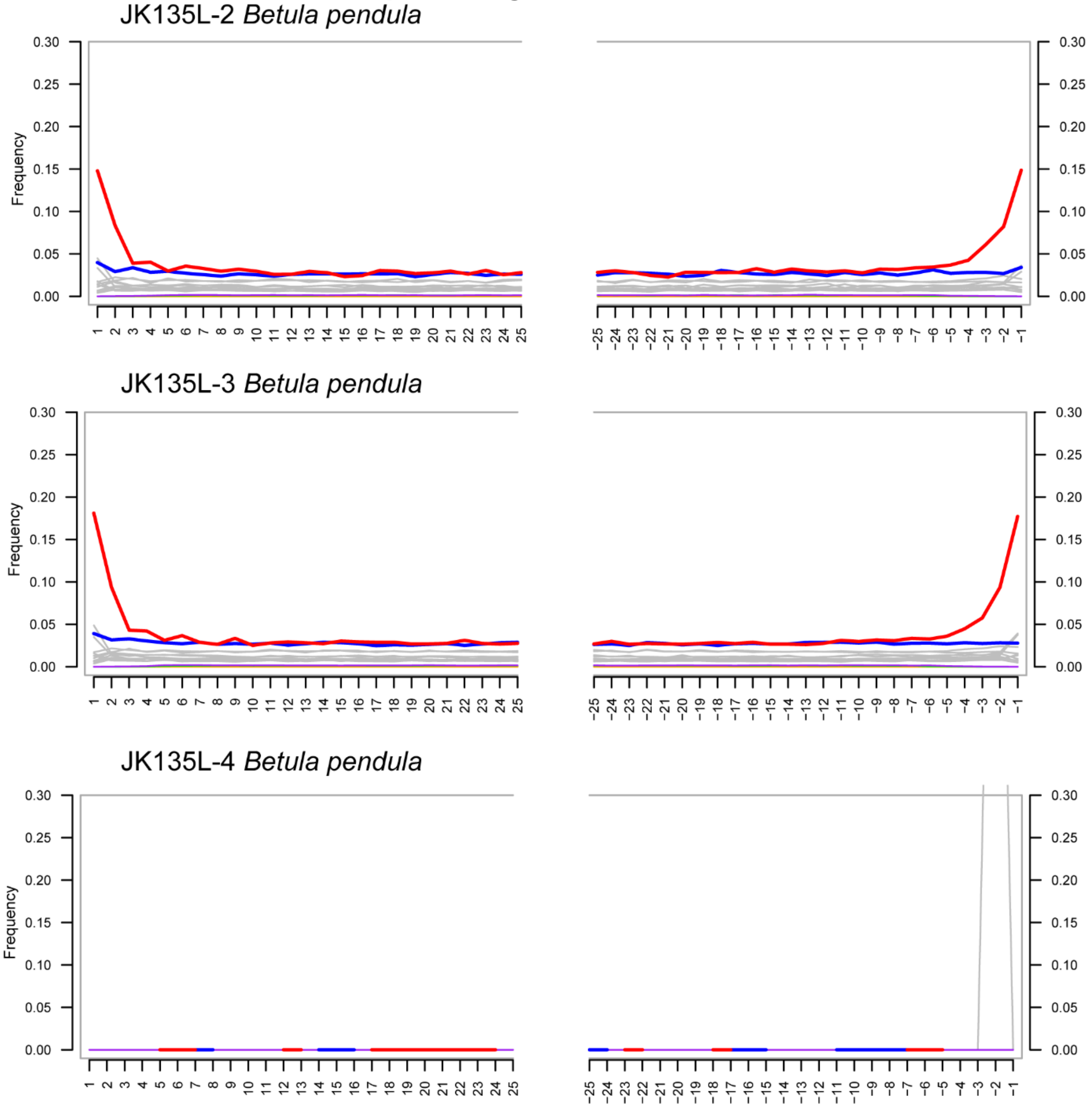

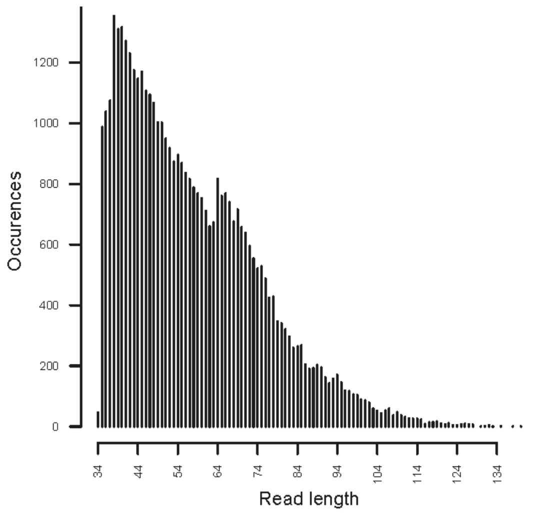

JK135L- 2 Betula pendula

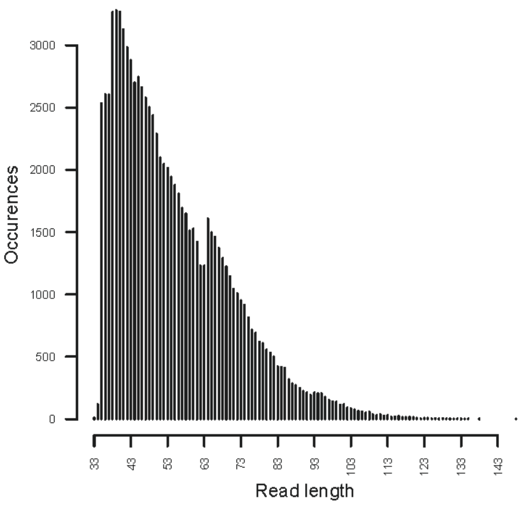

JK135L- 3 Betula pendula

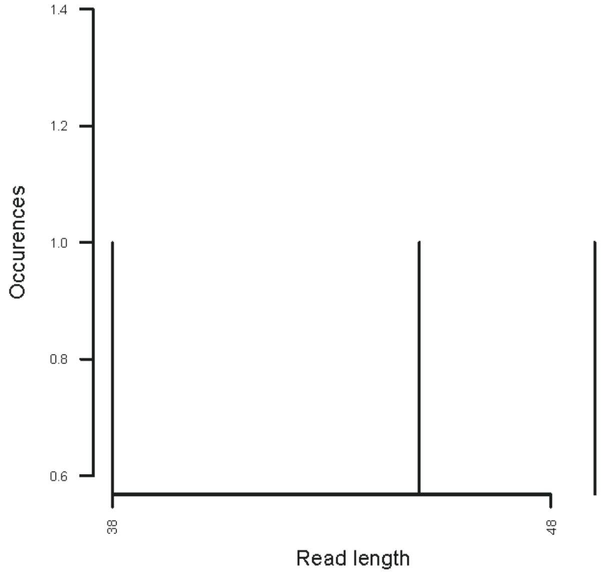

JK135L- 4 Betula pendula

Figure S2 *Betula pendula* *sed*aDNA read length and damage plots. JK135L-4 is the negative control. Damage patterns are consistent with ancient DNA. Plots generated by MapDamage.

**Figure S3- Read length distributions and ancient DNA deamination plots for reads mapped to the *Corylus avellana* genome**

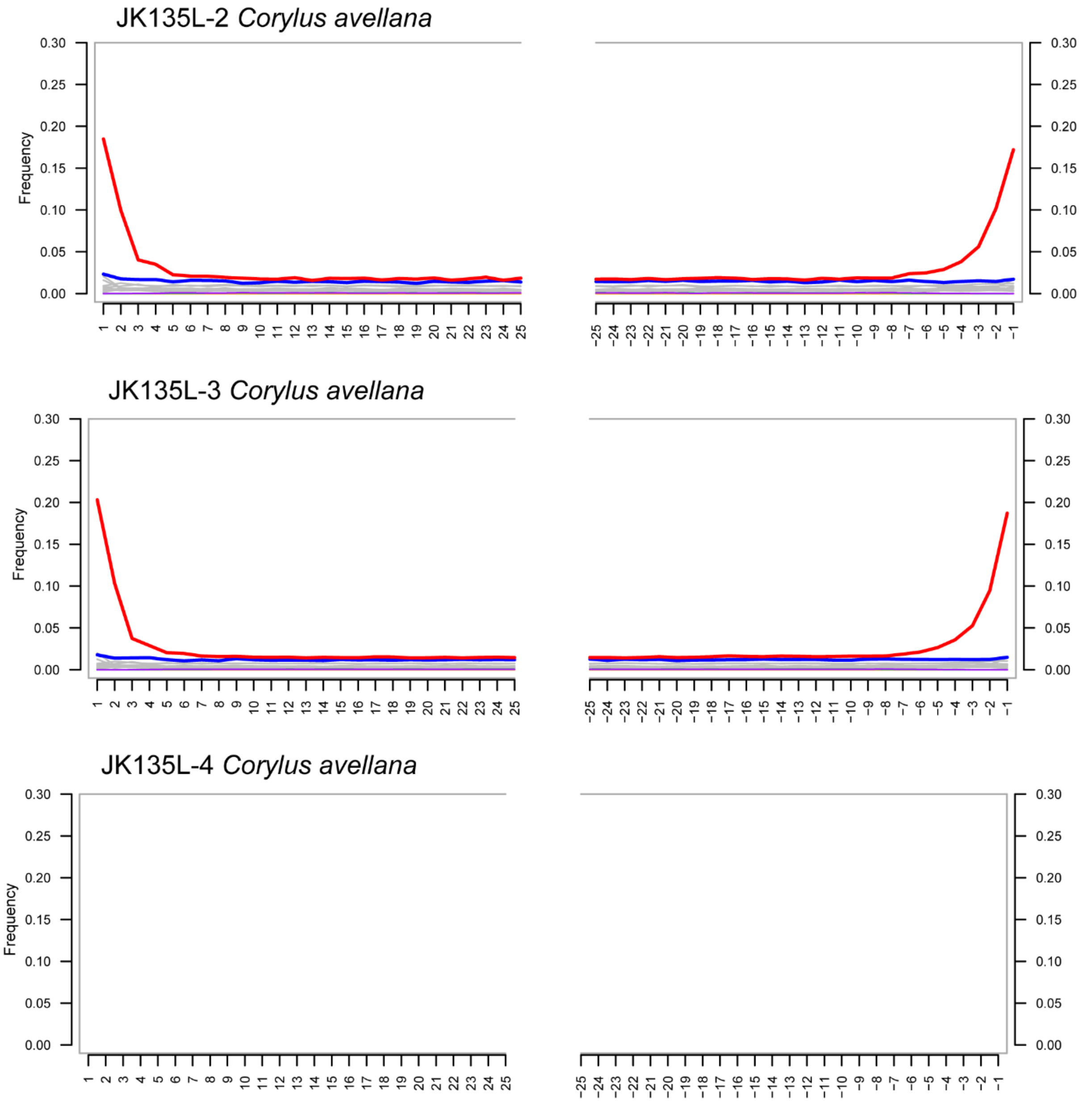

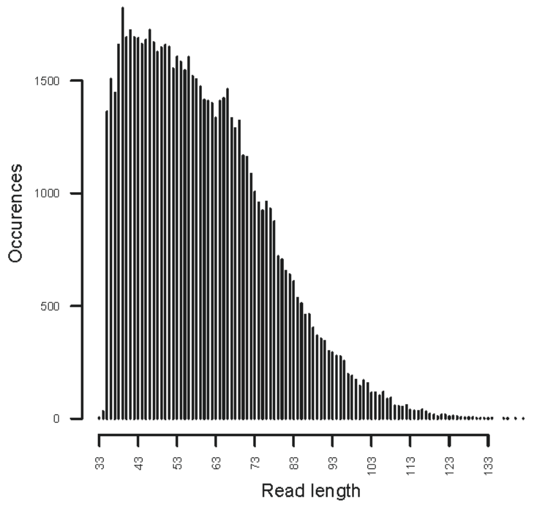

JK135L- 2 Corylus avellana

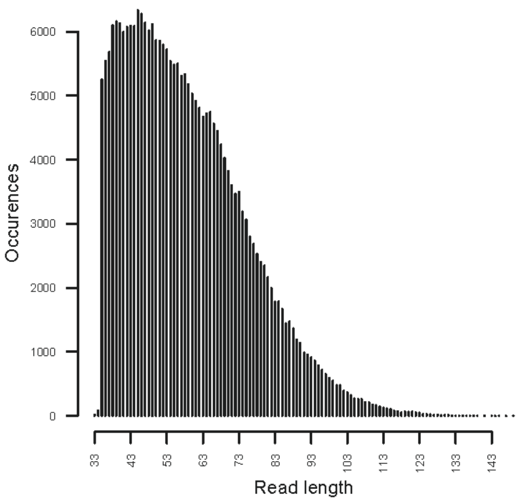

JK135L- 3 Corylus avellana

Figure S3 *Corylus avellana* *sed*aDNA read length and damage plots. JK135L-4 is the negative control. No reads were present in the negative control. Damage patterns are consistent with ancient DNA. Plots generated by MapDamage.

**Figure S4- Read length distributions and ancient DNA deamination plots for reads mapped to the *Salix brachista* genome**

JK135L-2 *Salix*

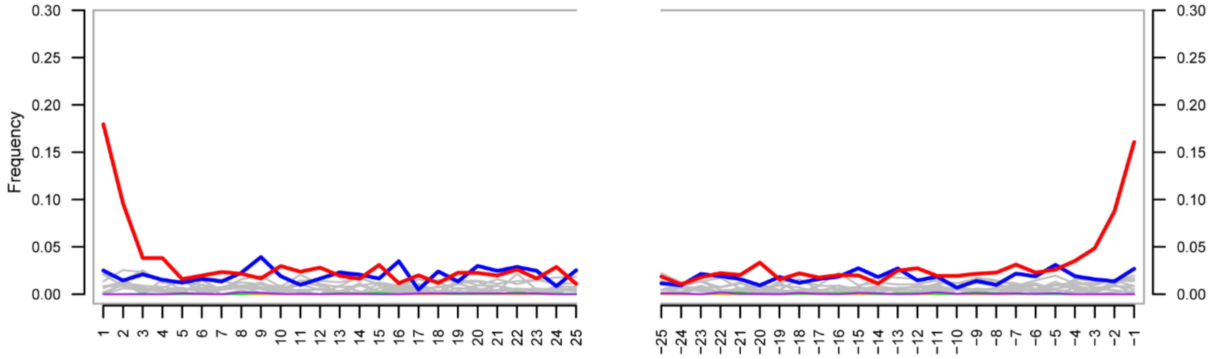

JK135L-3 *Salix*

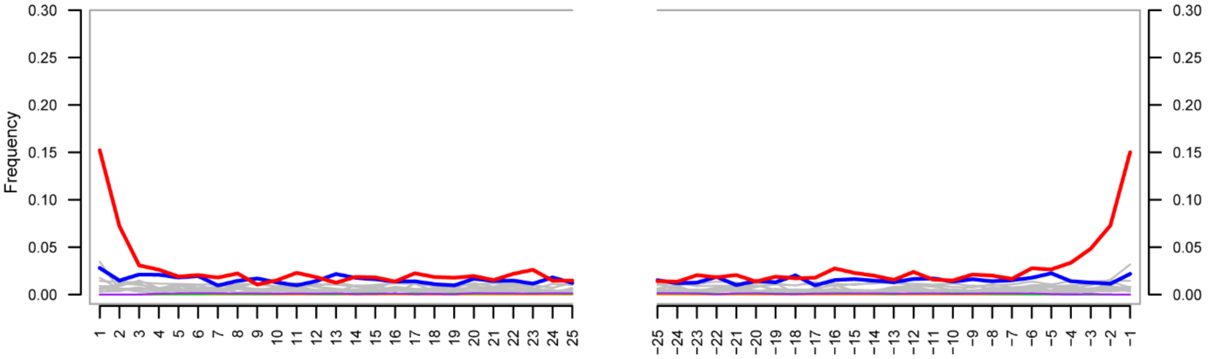

JK135L-4 *Salix*

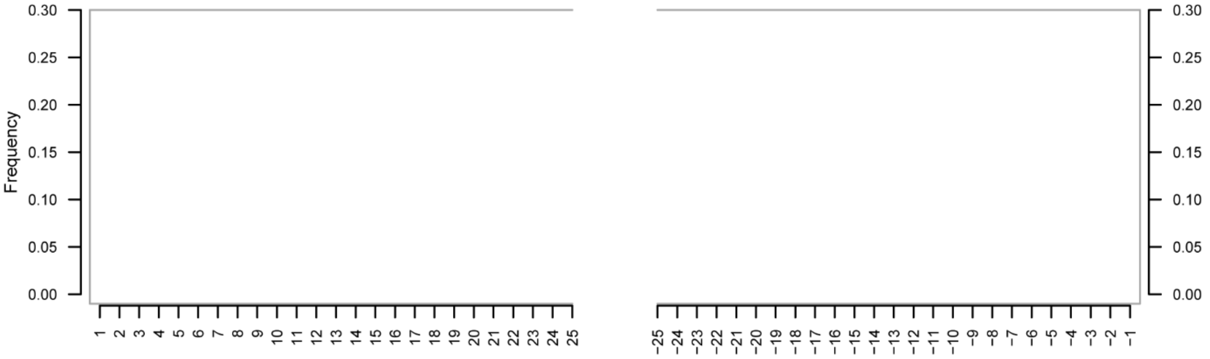

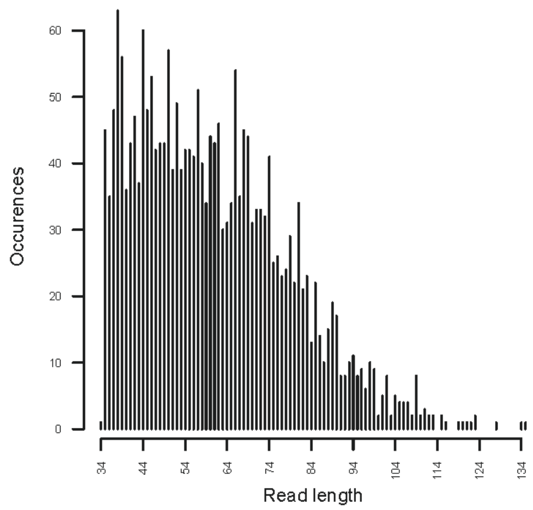

JK135L- 2 Salix

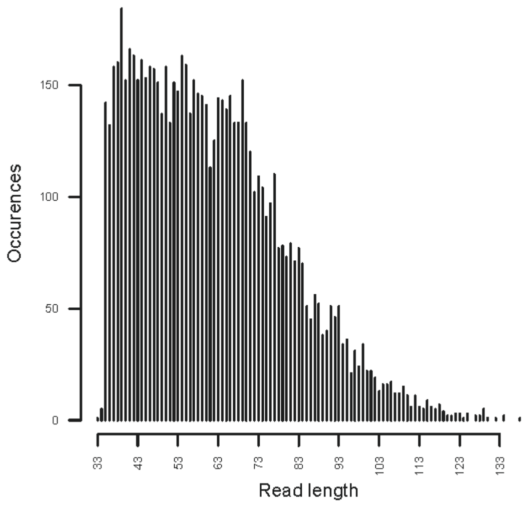

JK135L- 3 Salix

Figure S4 *Salix sed*aDNA read length and damage plots. JK135L-4 is the negative control. No reads were present in the negative control. Damage patterns are consistent with ancient DNA. Plots generated by MapDamage.

**Figure S5- Read length distributions and ancient DNA deamination plots for reads mapped to the *Homo sapiens* genome**

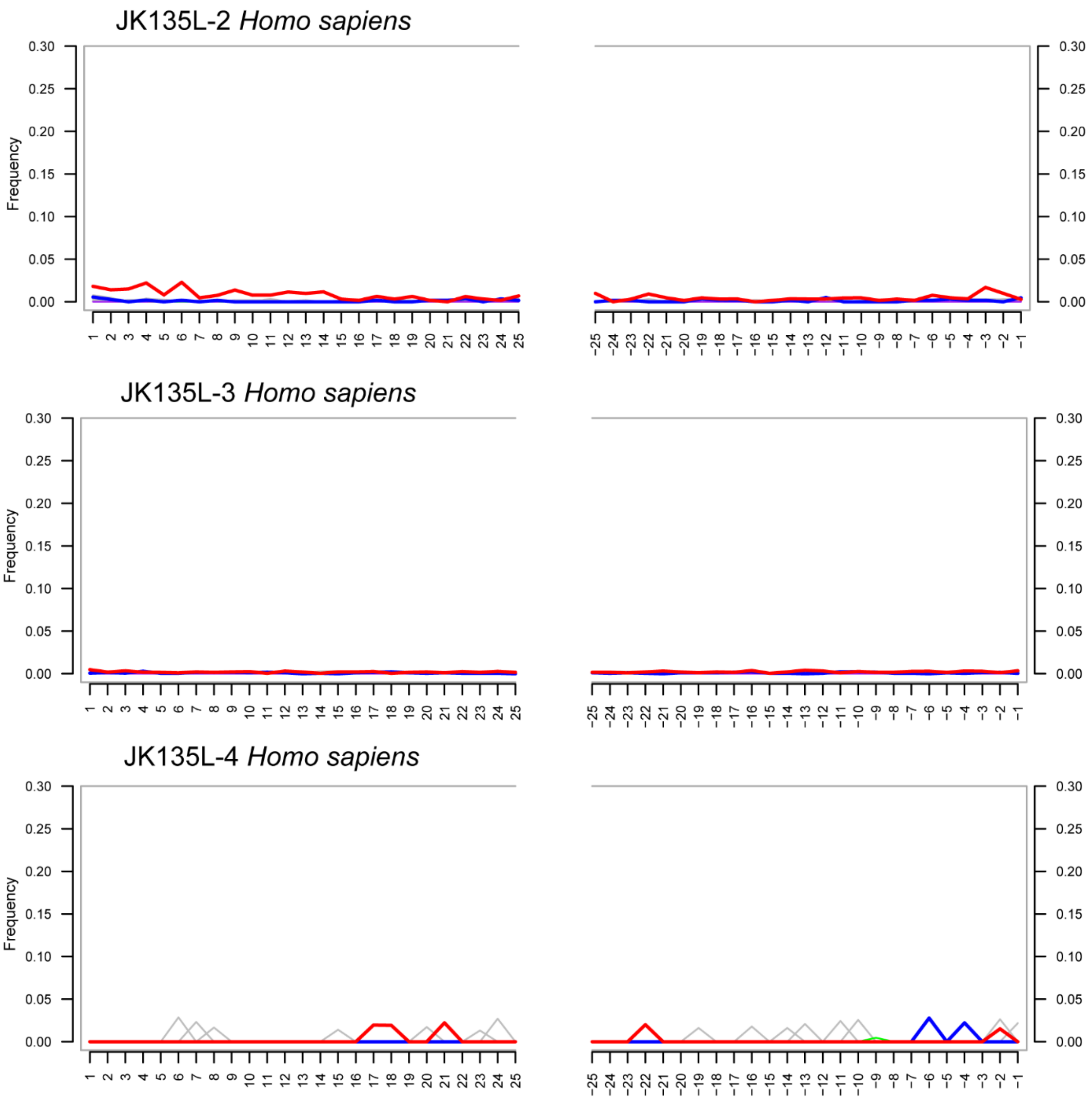

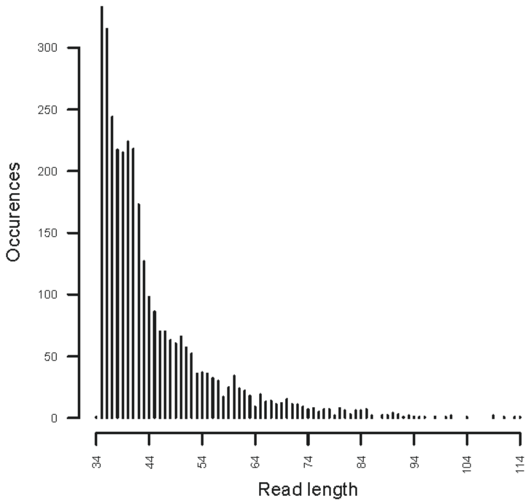

JK135L- 2 Homo sapiens

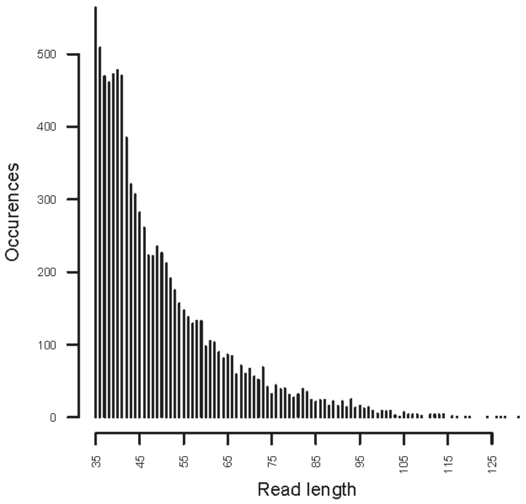

JK135L- 3 Homo sapiens

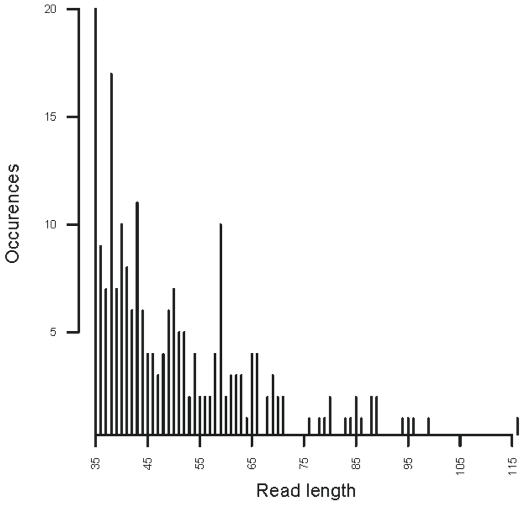

JK135L- 4 Homo sapiens

Figure S5 *Homo sapiens sed*aDNA read length and damage plots. JK135L-4 is the negative control. Plots generated by MapDamage.

**Figure S6- Read length distributions and ancient DNA deamination plots for reads mapped to the *Bos taurus* genome**

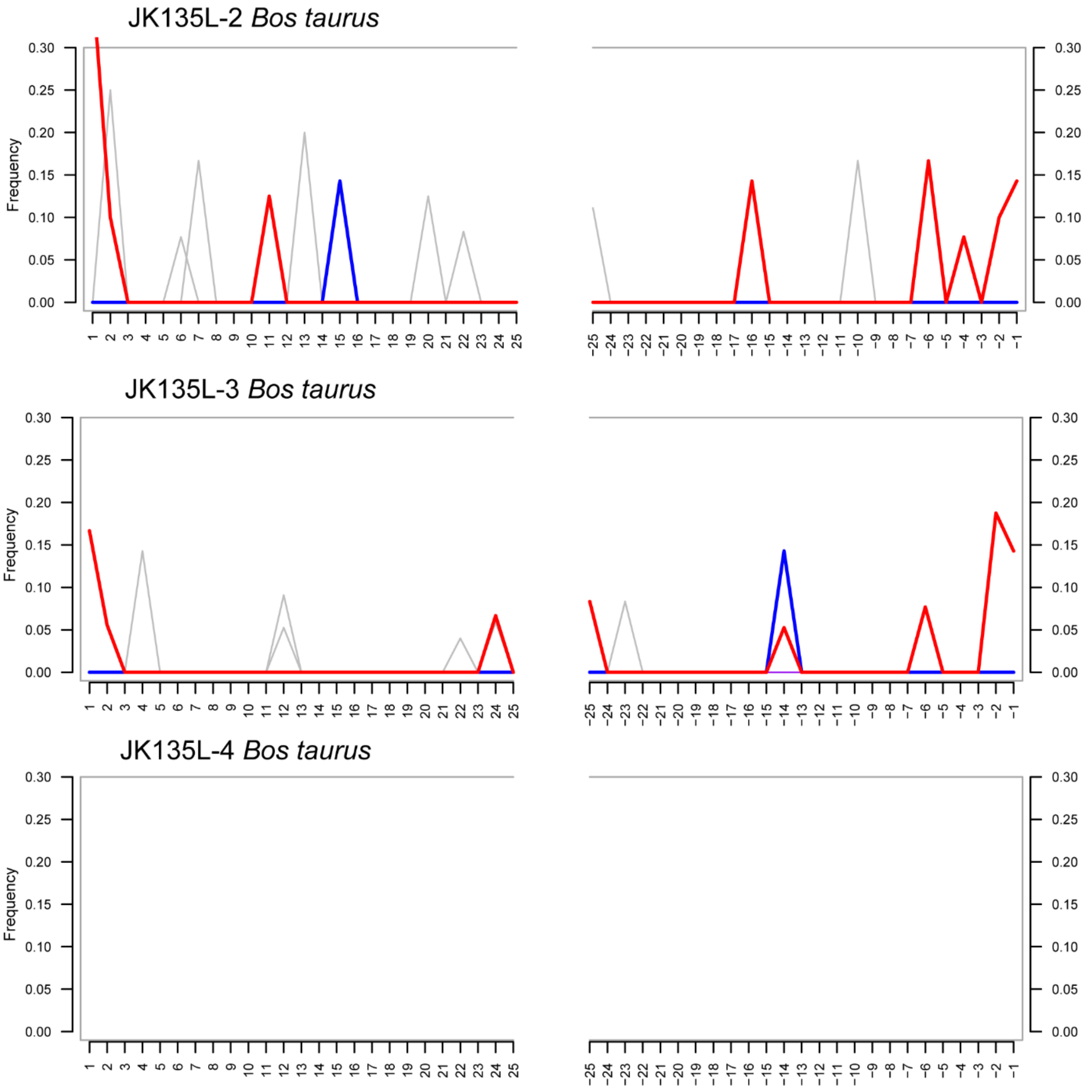

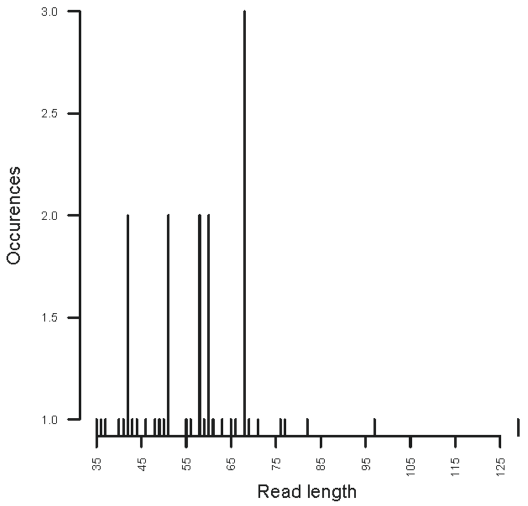

JK135L- 2 Bos taurus

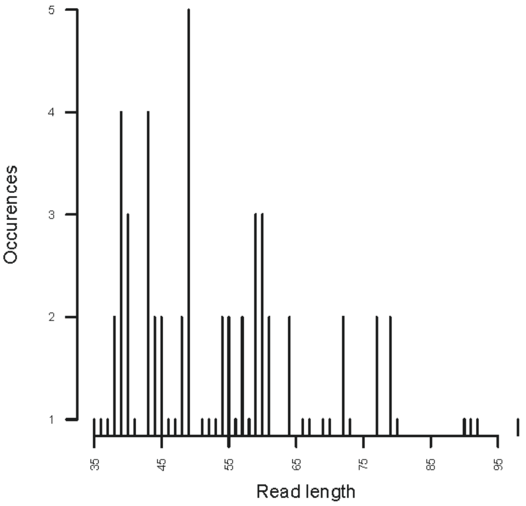

JK135L- 3 Bos taurus

Figure S6 *Bos taurus sed*aDNA read length and damage plots. JK135L-4 is the negative control. No reads were present in the negative control. The other two samples show damage consistent with ancient DNA. Plots generated by MapDamage.

### Figure S7- Read length distributions and ancient DNA deamination plots for reads mapped to the *Ovis aries* genome

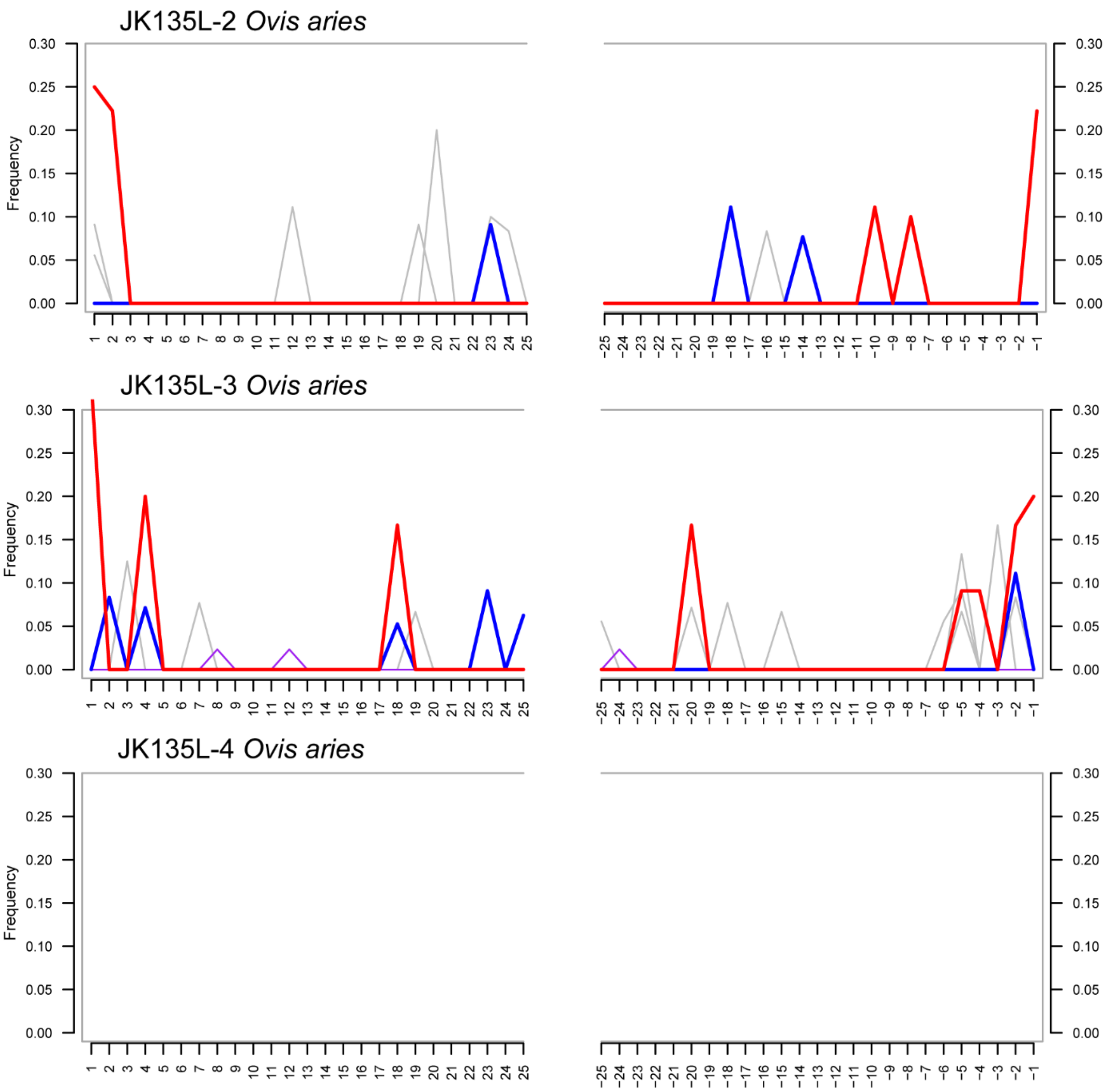

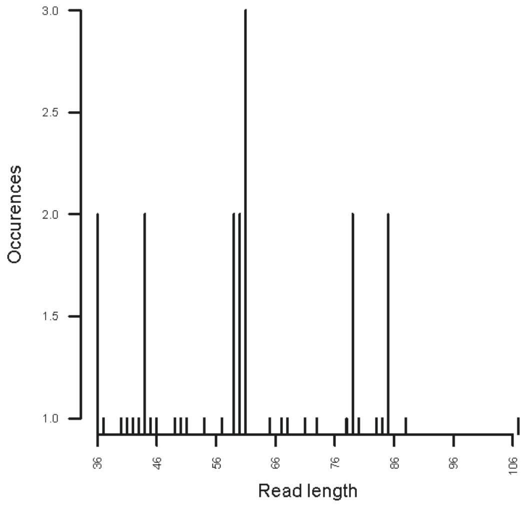

JK135L- 2 Ovis aries

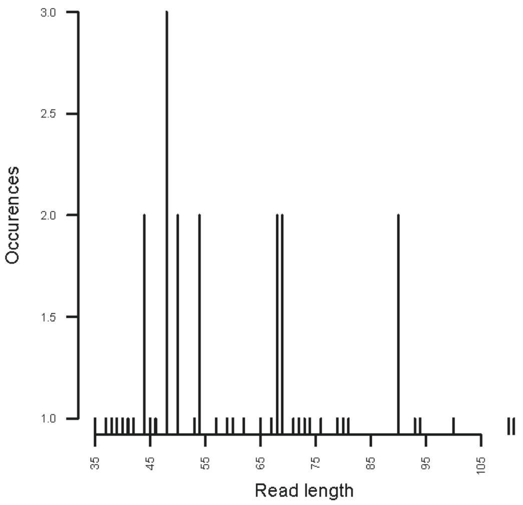

JK135L- 3 Ovis aries

Figure S7 *Ovis aries sed*aDNA read length and damage plots. JK135L-4 is the negative control. No reads were present in the negative control. Damage patterns are consistent with ancient DNA. Plots generated by MapDamage.

### Supplementary Material Reference List

Achilli, A., Bonfiglio, S., Olivieri, A., Malusa, A., Pala, M., Kashani, B. H., Torroni, A. *et al.* (2009). The multifaceted origin of taurine cattle reflected by the mitochondrial genome. *PloS one*, *4*(6), e5753.

Alsos, I. G., Lammers, Y., Kjellman, S. E., Merkel, M. K. F., Bender, E. M., Rouillard, A., & Schomacker, A. (2021). Ancient sedimentary DNA shows rapid post-glacial colonisation of Iceland followed by relatively stable vegetation until the Norse settlement (Landnám) AD 870. *Quaternary Science Reviews*, *259*, 106903.

Alsos, I. G., Lammers, Y., Yoccoz, N. G., Jørgensen, T., Sjögren, P., Gielly, L., & Edwards, M. E. (2018). Plant DNA metabarcoding of lake sediments: How does it represent the contemporary vegetation. *PloS one*, *13*(4), e0195403.

Alsos, I. G., Sjögren, P., Brown, A. G., Gielly, L., Merkel, M. K. F., Paus, A., & Van Der Bilt, W. G. (2020). Last Glacial Maximum environmental conditions at Andøya, northern Norway; evidence for a northern ice-edge ecological "hotspot". *Quaternary Science Reviews*, *239*, 106364.

Alsos, I. G., Rijal, D. P., Ehrich, D., Karger, D. N., Yoccoz, N. G., Heintzman, P. D., PhyloNorway Consortium. *et al.* (2022). Postglacial species arrival and diversity buildup of northern ecosystems took millennia. *Science advances*, *8*(39), eabo7434.

Borry, M., Hübner, A., Rohrlach, A. B. & Warinner, C. (2021) PyDamage: automated ancient damage identification and estimation for contigs in ancient DNA *de novo* assembly. *PeerJ*, *9*, e11845.

Boyer, F., Mercier, C., Bonin, A., Le Bras, Y., Taberlet, P., & Coissac, E. (2016). obitools: A unix-inspired software package for DNA metabarcoding. *Molecular ecology resources*, *16*(1), 176-182.

Camacho, C., Coulouris, G., Avagyan, V., Ma, N., Papadopoulos, J., Bealer, K., & Madden, T. L. (2009). BLAST+: architecture and applications. *BMC bioinformatics*, *10*(1), 1-9.

Chen, W., & Ficetola, G. F. (2020). Numerical methods for sedimentary-ancient-DNA-based study on past biodiversity and ecosystem functioning. *Environmental DNA*, *2*(2), 115-129.

Feuerborn, T. R., Palkopoulou, E., van der Valk, T., von Seth, J., Munters, A. R., Pečnerová, P., Díez-del-Molino, D. *et al.* (2020). Competitive mapping allows for the identification and exclusion of human DNA contamination in ancient faunal genomic datasets. *BMC genomics*, *21*(1), 1-10.

Fonseca, V. G. (2018). Pitfalls in relative abundance estimation using eDNA metabarcoding.

Gansauge, M. T., & Meyer, M. (2013). Single-stranded DNA library preparation for the sequencing of ancient or damaged DNA. *Nature protocols*, *8*(4), 737-748.

Gansauge, M. T., Gerber, T., Glocke, I., Korlević, P., Lippik, L., Nagel, S., Meyer, M. *et al.* (2017). Single-stranded DNA library preparation from highly degraded DNA using T4 DNA ligase. *Nucleic acids research*, *45*(10), e79-e79.

Garcés-Pastor, S., Coissac, E., Lavergne, S., Schwörer, C., Theurillat, J. P., Heintzman, P. D., Alsos, I. G. *et al.* (2022) High resolution ancient sedimentary DNA shows that alpine plant diversity is associated with human land use and climate change. *Nature communications*, *13*(1), 6559.

Giguet-Covex, C., Pansu, J., Arnaud, F., Rey, P. J., Griggo, C., Gielly, L., Taberlet, P. *et al.* (2014). Long livestock farming history and human landscape shaping revealed by lake sediment DNA. *Nature communications*, *5*(1), 3211.

Heiri, O., Lotter, A. F., & Lemcke, G. (2001). Loss on ignition as a method for estimating organic and carbonate content in sediments: reproducibility and comparability of results. *Journal of paleolimnology*, *25*(1), 101-110.

Hudson, S. M., Pears, B., Jacques, D., Fonville, T., Hughes, P., Alsos, I., Brown, A. *et al.* (2022). Life before Stonehenge: The hunter-gatherer occupation and environment of Blick Mead revealed by sedaDNA, pollen and spores. *Plos one*, *17*(4), e0266789.

Jónsson, H., Ginolhac, A., Schubert, M., Johnson, P. L., & Orlando, L. (2013). mapDamage2. 0: fast approximate Bayesian estimates of ancient DNA damage parameters. *Bioinformatics*, *29*(13), 1682-1684.

Li, H., & Durbin, R. (2010). Fast and accurate long-read alignment with Burrows–Wheeler transform. *Bioinformatics*, *26*(5), 589-595.

Li, H., Handsaker, B., Wysoker, A., Fennell, T., Ruan, J., Homer, N., Durbin, R. *et al.* (2009). The sequence alignment/map format and SAMtools. *Bioinformatics*, *25*(16), 2078-2079.

Peltzer, A., Jäger, G., Herbig, A., Seitz, A., Kniep, C., Krause, J., & Nieselt, K. (2016). EAGER: efficient ancient genome reconstruction. *Genome biology*, *17*(1), 1-14.

Rossi, C., Sinding, M.-H. S., Mullin, V. E., Scheu, A., Erven, J. A. M., Verdugo, M. P., Daly, K. G. *et al*. (2024) The genomic natural history of the aurochs. *Nature*, *635*, 136-141.

Schubert, M., Ginolhac, A., Lindgreen, S., Thompson, J. F., Al-Rasheid, K. A., Willerslev, E., Orlando, L. *et al.* (2012). Improving ancient DNA read mapping against modern reference genomes. *BMC genomics*, *13*(1), 1-15.

Schulte, L., Bernhardt, N., Stoof-Leichsenring, K., Zimmermann, H. H., Pestryakova, L. A., Epp, L. S., & Herzschuh, U. (2021). Hybridization capture of larch (Larix Mill.) chloroplast genomes from sedimentary ancient DNA reveals past changes of Siberian forest. *Molecular Ecology Resources*, *21*(3), 801-815.

Shen, W., & Ren, H. (2021). TaxonKit: A practical and efficient NCBI taxonomy toolkit. *Journal of Genetics and Genomics*, *48*(9), 844-850.

Simpson, J. T., & Durbin, R. (2012). Efficient de novo assembly of large genomes using compressed data structures. *Genome research*, *22*(3), 549-556.

Soininen, E. M., Gauthier, G., Bilodeau, F., Berteaux, D., Gielly, L., Taberlet, P., Yoccoz, N. G *et al.*. (2015). Highly overlapping winter diet in two sympatric lemming species revealed by DNA metabarcoding. *PloS one*, *10*(1), e0115335.

Sønstebø, J. H., Gielly, L., Brysting, A. K., Elven, R., Edwards, M., Haile, J., Brochmann, C *et al*. (2010). Using next-generation sequencing for molecular reconstruction of past Arctic vegetation and climate. *Molecular Ecology Resources*, *10*(6), 1009-1018.

Valentini, A., Taberlet, P., Miaud, C., Civade, R., Herder, J., Thomsen, P. F., Dejean, T. *et al.* (2016). Next-generation monitoring of aquatic biodiversity using environmental DNA metabarcoding. *Molecular ecology*, *25*(4), 929-942.

Willerslev, E., Davison, J., Moora, M., Zobel, M., Coissac, E., Edwards, M. E., Taberlet, P *et al*. (2014). Fifty thousand years of Arctic vegetation and megafaunal diet. *Nature*, *506*(7486), 47-51.
